# Is Probiotic Supplementation During Pregnancy Safe for Healthy Populations? A Systematic Review of Preclinical Studies

**DOI:** 10.1101/2024.11.04.621894

**Authors:** Mario R. Coca, Cristian Perez-Fernandez, Diego Ruiz-Sobremazas, Miguel Morales-Navas, Teresa Colomina, Caridad López-Granero, Fernando Sanchez-Santed

**Author notes:** **Corresponding authors**: Perez-Fernandez, Cristian & Sánchez- Santed, Fernando. **First author**: Coca, Mario R.

## Abstract

Gut-brain axis and probiotics supplementation during gestational period have gained an especial attention in the field of neurodevelopmental disorders. However, the influence of prenatal probiotics in physiology and neurodevelopment of healthy organisms is widely unexplored and probiotics are commercially such as dietetic supplementations without the necessity of ensuring safety in healthy organisms, as is required for drugs. The aim is to present and discuss the various biochemical, developmental and behavioral effects that prenatal probiotics have on the offspring of healthy organisms in rodent models and assess their security during gestation. We included a total of 15 preclinical studies discarding all those that did not have an appropriate control for the probiotic-treated group. Results suggest that probiotics, particularly *Lactobacillus* strains, influenced anxiety-like behaviors in mice and GABAergic system maturation, showing an apparent biphasic effect, reducing GABA functioning in early life but increasing them in later stages; and both prenatal *Lactobacillus* and *Bifidobacterium*, influenced microbiota and general development of healthy organisms. However, the significant variations in methodologies used across studies complicate the ability to elucidate specific conclusions related to bacterial strains or doses. Further research is imperative to ensure the safety and regulation of probiotic interventions, particularly during the gestational period.

## 1. INTRODUCTION

In recent decades, our understanding of the significant long-term impact of the prenatal environment on health has grown considerably, with particular emphasis on neurodevelopmental disorders (Doi et al., 2022). In this context, the gut microbiota and its connection to the formation, development, and proper functioning of the nervous system have garnered considerable attention within the field of basic neuroscience and in proposing interventions (Yan et al., 2023). Alterations in gut microbiota have been shown to impact brain physiology and behavior through mechanisms like direct vagus nerve activation, immune system interactions, and the production of microbial metabolites such as short-chain fatty acids (SCFAs), neurotransmitters or its precursors (Sampson and Mazmanian, 2015; Sherwin et al., 2019). The gut microbiome begins developing at birth and is influenced by factors like delivery method, diet, and age.

While it stabilizes around age 3, it continues to be shaped by hormonal and environmental changes. In older adults, particularly after 65, microbial diversity decreases significantly, and there is a high degree of individual variation in gut microbiome composition (Bicknell et al., 2023).

The disruption of the microbiota balance is known as gut dysbiosis. Dysbiosis has been linked to neurodevelopmental and neurodegenerative disorders by the generating of disturbances such as increased blood-brain barrier permeability, altering the circulating levels of cytokines and other immune-related molecules, corticotrophin levels, kidney and cardiovascular health, serotonin and mental health, gut barrier damage, altering the general metabolic status and microglia activation (Bicknell et al., 2023). In this context, therapeutic probiotic interventions —live microorganisms, primarily bacteria, that confer health benefits to their host when consumed in adequate amounts— (Fijan, 2014) have received much attention in research (Sanders et al., 2013). Probiotics can affect the development of offspring either by colonizing the fetus through the placenta and amniotic fluid or by altering fetal growth indirectly through changes in maternal health and metabolism (Swartwout and Luo, 2018). In this regard, probiotic interventions in pregnant women show potential in reducing gestational diabetes risk, improving inflammatory responses, and managing blood pressure and insulin sensitivity, though effects on glycemic control vary across strains (Abbasi et al., 2021). In preclinical literature, a non-systematic review conducted by Cuinat et al. (2022), showed that gestational probiotic supplementation in animal models influence offspring outcomes.

These effects include reductions in body weight, changes in digestive enzyme activity, gut structure and function, modifications in lipid profiles, enhanced levels of IgA and anti-inflammatory cytokines, shifts in beneficial gut microbiota, improvements in insulin sensitivity and fat mass and increase in cortical neuron density and behavioral positive changes related to anxiety (Cuinat et al., 2022). However, they analyzed the effects of gestational supplementation in animal pathological models rather than in healthy control organisms.

Since it is considered beneficial for patients and likely harmless or even beneficial for normative individuals, an increasing number of people without diagnosed conditions are using these substances. However, little is known about their effects on normative groups, as this has generally not been a focus of scientific research. In recent years, a growing body of literature has emerged that questions the assumed safety attributed to the use of probiotics, especially during pregnancy. Theoretically, it has been proposed that probiotics may have detrimental effects on hosts, potentially leading to systemic infections, disruptions in neurodevelopment, changes in metabolic functions, excessive immune activation, effects on metabolism, and alterations in drug responses and toxicity (Kolaček et al., 2017; Merenstein et al., 2023; Doron and Snydman, 2015).

Because in most countries around the world, probiotics are considered dietary supplements or beneficial natural products, they do not undergo extensive safety testing or quality control, as is the case with drugs (Kolaček et al., 2017). In this world context, more security evidence in probiotic gestational interventions should be found, especially given the fact that prenatal period is the most critical timeframe for inducing neurodevelopmental alterations (Doi et al., 2022) and probiotics are often recommended during pregnancy to reduce the risk of several pathologies (Sanz, 2011; Tette et al., 2022), it is essential to ensure that their use does not inadvertently alter the physiological development of the offspring. This is why the objective of this systematic review is to investigate the outcomes of preclinical studies regarding the impact of prenatal probiotic supplementations on health control rodent models, assessing safety considerations across various behavioral, developmental, and biochemical parameters in the short, medium and long term.

## 2. METHODOLOGY

### 2.1 Search strategy

The bibliographical search for this systematic review was completed in December 2023 and it is based on the Preferred Reporting Items for Systematic Reviews and Meta- Analyses (PRISMA) Statement (Shamseer et al., 2015) and the practical guide for systematic reviews in health sciences (Cajal et al., 2020). In order to search and find studies to date relevant to the objective set, the following databases were used: Web of Science, Scopus and PUBMED. The keywords and search equation used were “probiotic” AND “brain” OR “neuro*” AND “prenatal” OR “gestational” OR “maternal” AND “behavior” OR “emotion” OR “anxi*” OR “learning” OR “memory” OR “cognition” OR “sociability” OR “impulsivity” OR “compulsivity” AND “rat” OR “mouse” OR “mice”. A second search was conducted in September 2024 to include new studies published after the initial search. The same databases and search equations were used, with the additional filter of the year “2024” to capture the most recent research prior to finalizing the systematic review for publication. Both searches will be reflected together in the results section.

### 2.2 Elegibility criteria

The purpose of this review is to shed light on the potential short-term, medium-term, and long-term effects that the intake of probiotic supplements during pregnancy may have on offspring, in studies involving rodent animal models. Human and other animal models were excluded. All prenatal probiotic supplementation should include exposures that cover the gestational period, even if they extend into the postnatal period. Based on this, we excluded studies that involved chronic treatments extending into adulthood or only postnatal treatments. The key criterion for the inclusion of such interventions is that they are conducted on healthy control subjects. Effects on neurodevelopmental models or subjects exposed to toxics will not be considered. Moreover, we considered articles that incorporated vitamins or other dietary supplements alongside probiotics as excipients, given that they are commonly marketed together (Abboud et al., 2020).

There is no eligibility criterion for the publication year, as there are no similar previous systematic reviews in rodent control subjects. The primary outcomes to be assessed will include biochemical variables (such as genetic, epigenetic, protein, or metabolomic markers), microbiota composition changes, and quantitative behavioral or developmental measures at each age and developmental milestone of the offspring if supplementation was received during gestation.

### 2.3 Study selection and data extraction

We analyzed all identified articles to prevent bias, utilizing the IT tool Rayyan (Ouzzani et al., 2016). Any disagreements were discussed until a consensus was reached. We didn’t just limit our screening to titles and abstracts; each topic article was thoroughly read. This was necessary because effects on controls are often not fully captured in the abstract. Those studies that did not meet our criteria were excluded.

### 2.4 Quality assessment

To assess the risk of bias in animal studies, the SYRCLE tool was used to evaluate bias risk (Hooijmans et al., 2014), which is an adaptation of the RoB tool based on Cochrane guidelines. This method has also been used in other systematic reviews (Biosca-Brull et al., 2021). The SYRCLE tool consists of five quality domains: selection, performance, detection, attrition, and reporting bias. It assigns the following maximum points: six for selection, four for performance, four for detection, four for attrition, four for reporting bias, and two for other biases (total score of 26). Articles are categorized according to the total bias score: 0-8 (Low), 9-17 (Moderate), and 18-26 (High).

## 3. RESULTS

### 3.1 Selection of studies and bias assessment

A flow diagram illustrates the whole search strategy (Fig.1). The search yielded a total of 250 studies and, after excluded 74 duplicates a total of 176 articles were selected. Out of those 176 studies, 42 were excluded based on a review of titles and abstracts for being unrelated to the topic. In the second screening (n=134), we assessed main text and supplementary material. A total of 121 articles were excluded due to the following reasons: 43 focused on postnatal treatment, 21 lacked an appropriate control group for the probiotic supplementation, 22 were reviews, and 35 did not involve any probiotic treatment. A parallel search was conducted, which identified two additional articles that were included in the review. Thus, 10 mouse model studies and 5 rat model studies comprised the total number of studies included in this review. Of the 15 studies included, 6 focus exclusively on prenatal treatment, while the remaining 9 extend the treatment into the early postnatal period. Regarding bias assessment (see Tables 1 and 2), 66,7% of the articles included had low bias and 33,3% had medium bias. None of the articles in either section presented a high bias score.

**Figure 1.**
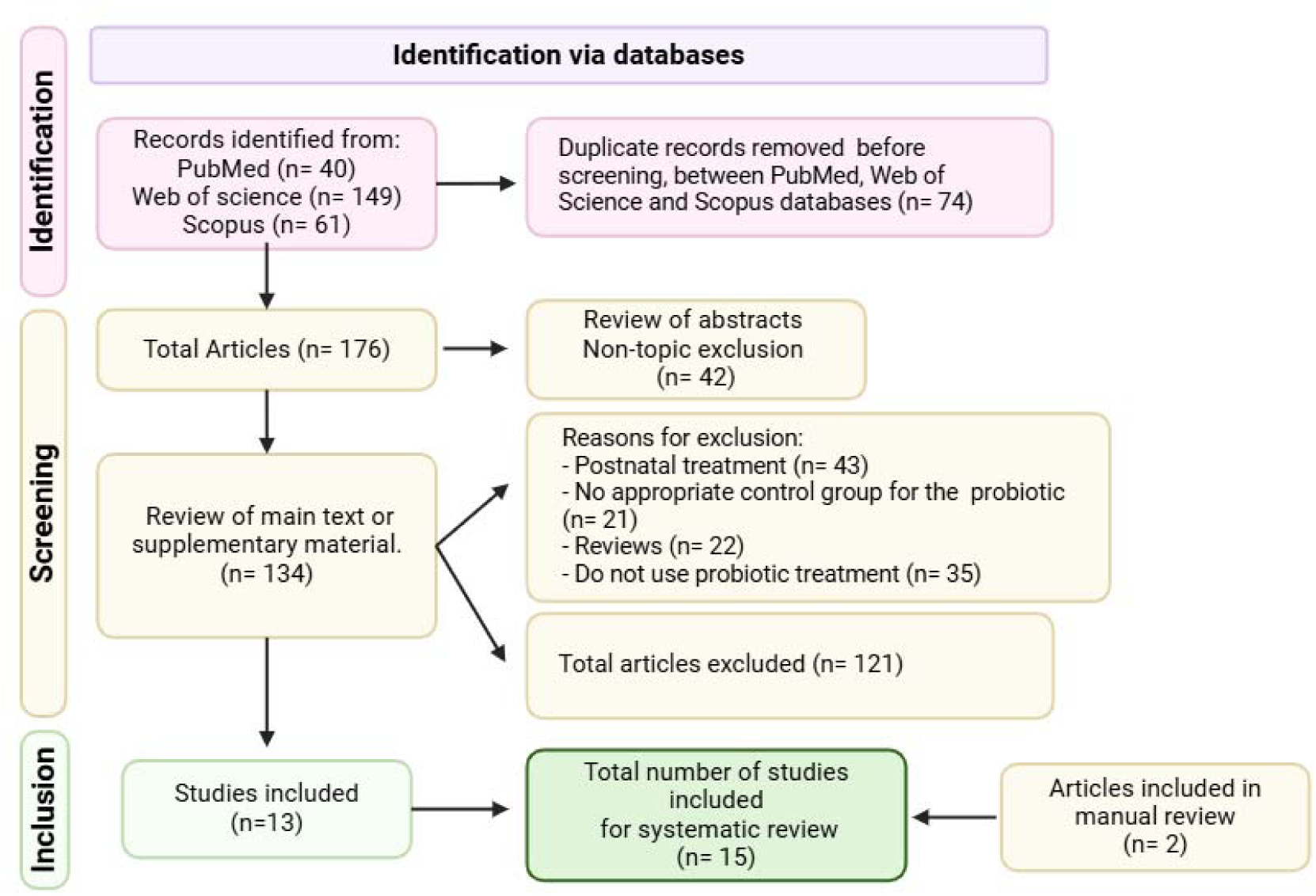
Search flow diagram

**Table 1.**
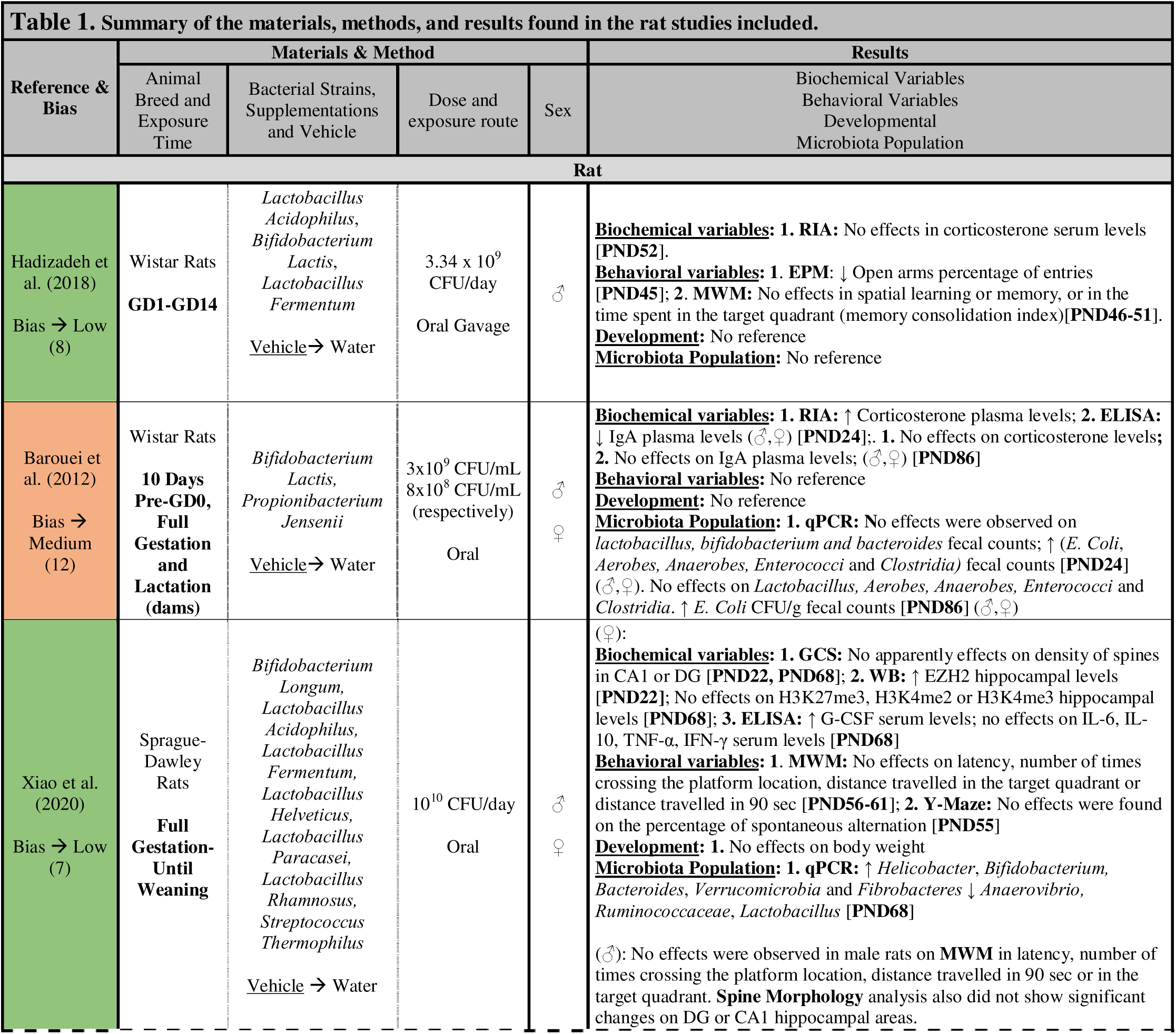
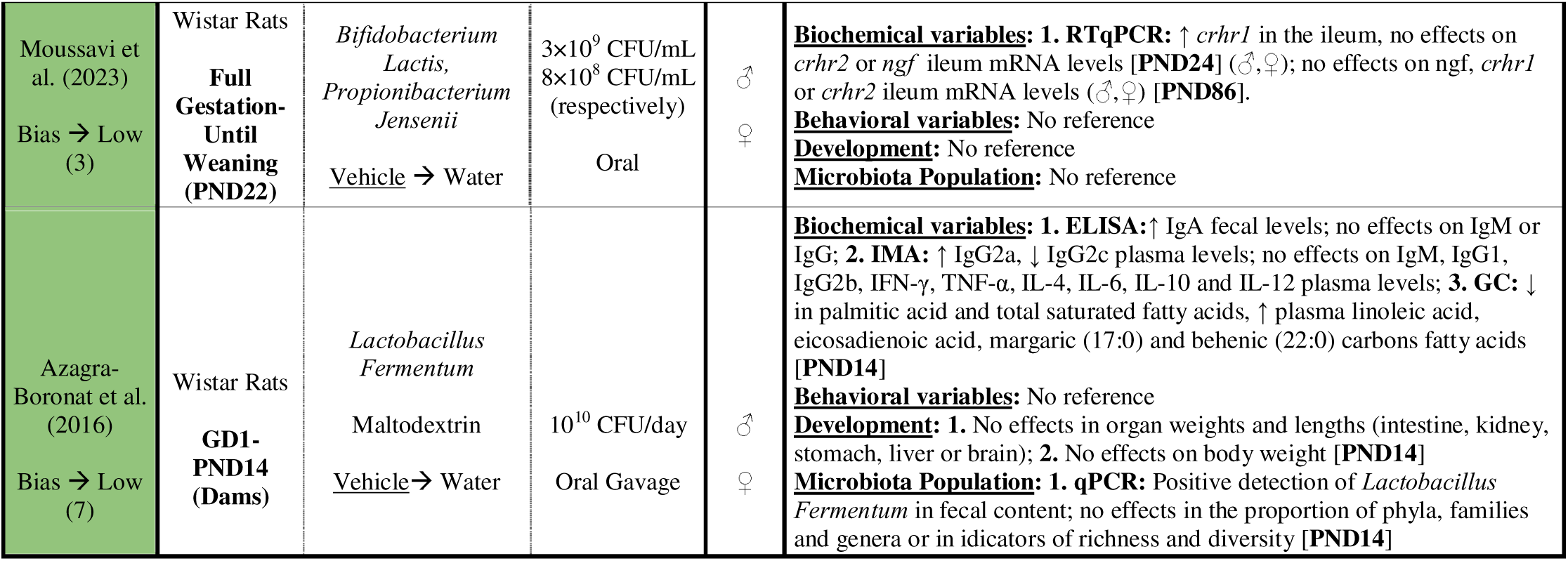
Summary of the materials, methods, and results found in the rat studies included.

Abbr. PND: Postnatal day; GD: Gestational day; HPAC: High-Performance Liquid 217 Chromatography; GC-MS: Gas Chromatography Mass Spectrometry; RIA: Radioimmunoassay; IMA: Immunology Multiplex Assay; LC-MS: Liquid Chromatography-Mass Spectrometry; IFS: Immunofluorescence Staining; HPLC: High-Performance Liquid Chromatography; ELISA: Enzyme-Linked ImmunoSorbent Assay; EIA: Non-ELISA Enzyme Immunoassay; DGGE: Denaturing Gradient Gel Electrophoresis; WB: Western Blotting; IHC: Immunohistochemistry; GCS: Golgi-Cox Staining; LDB: Light-Dark Box Test; FST: Forced Swim Test; MBT: Marble Burying Test; RSG: Repetitive Self-Grooming Test; MVF: Manual Von Frey Test; RTqPCR: Quantitative Reverse Transcription PCR; 3CT: Three-Chamber Test; EPM: Elevated Plus Maze; TST: Tail Suspension Test; OFT: Open Field Test; ROTAROD: RotaRod Performance Test; RT-PCRarray: Reverse Transcription PCR Array; PMT: Plus Maze Test; 16S rRNA seq: 16S Ribosomal RNA Sequencing.

**Table 2.**
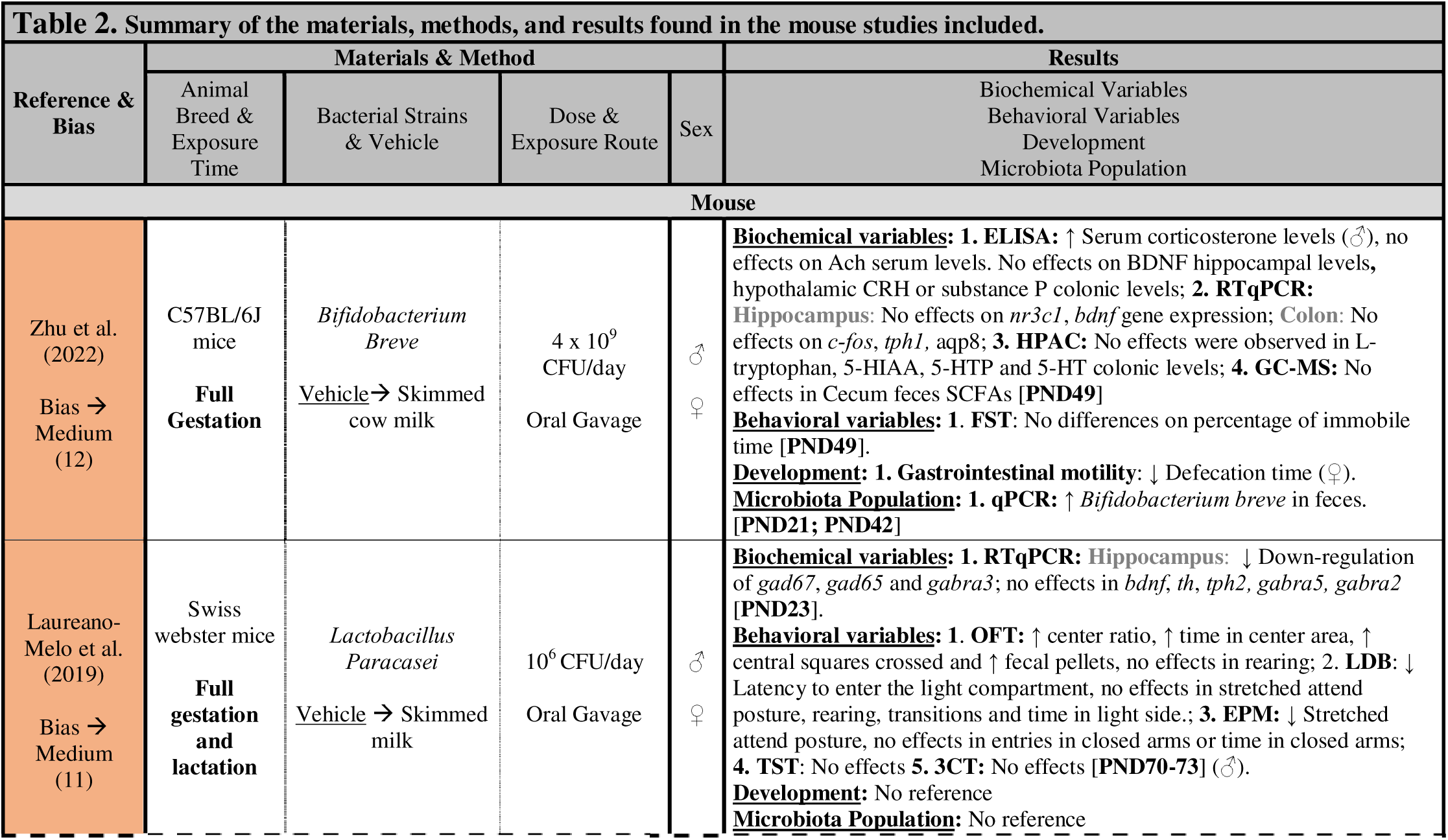
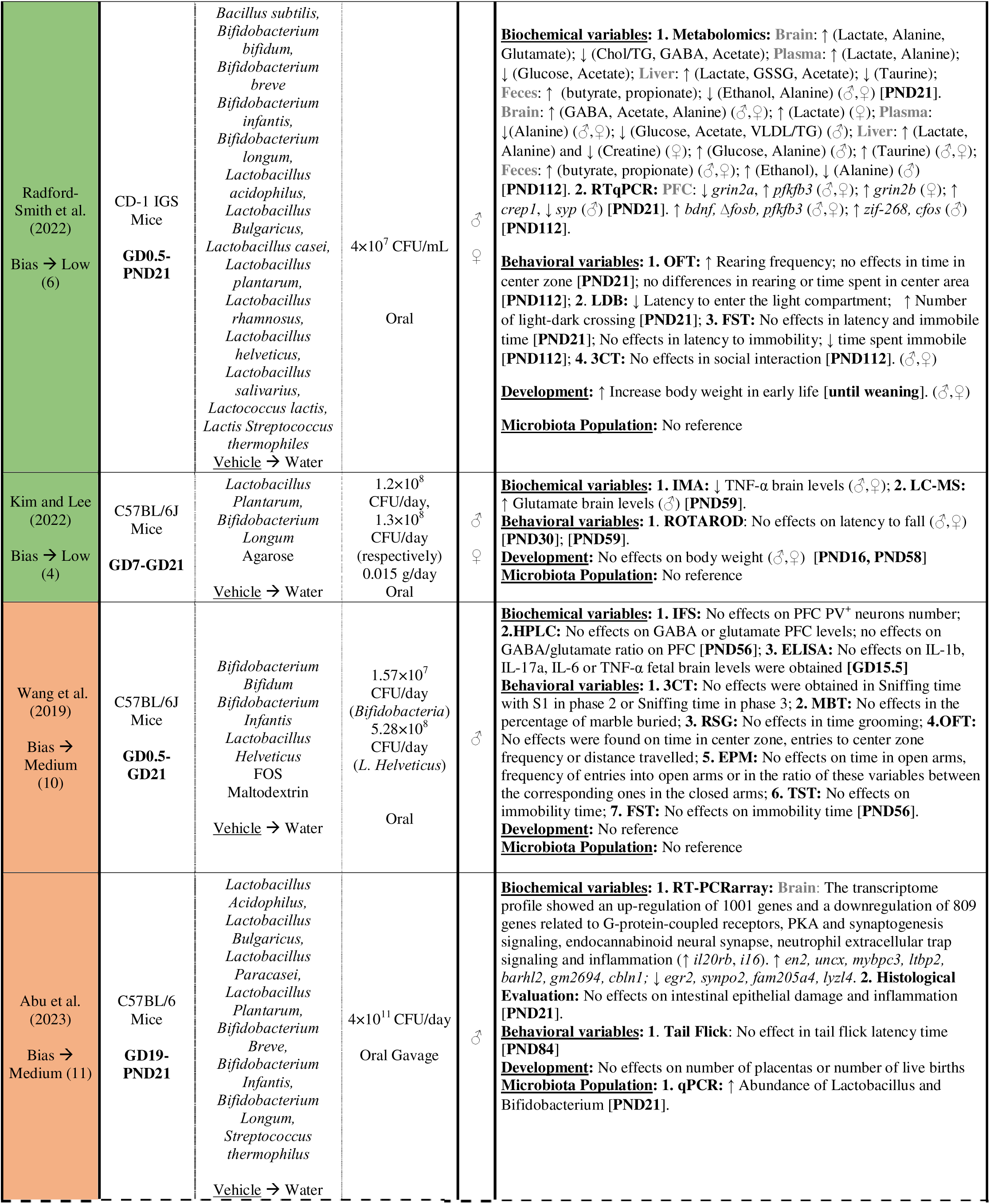
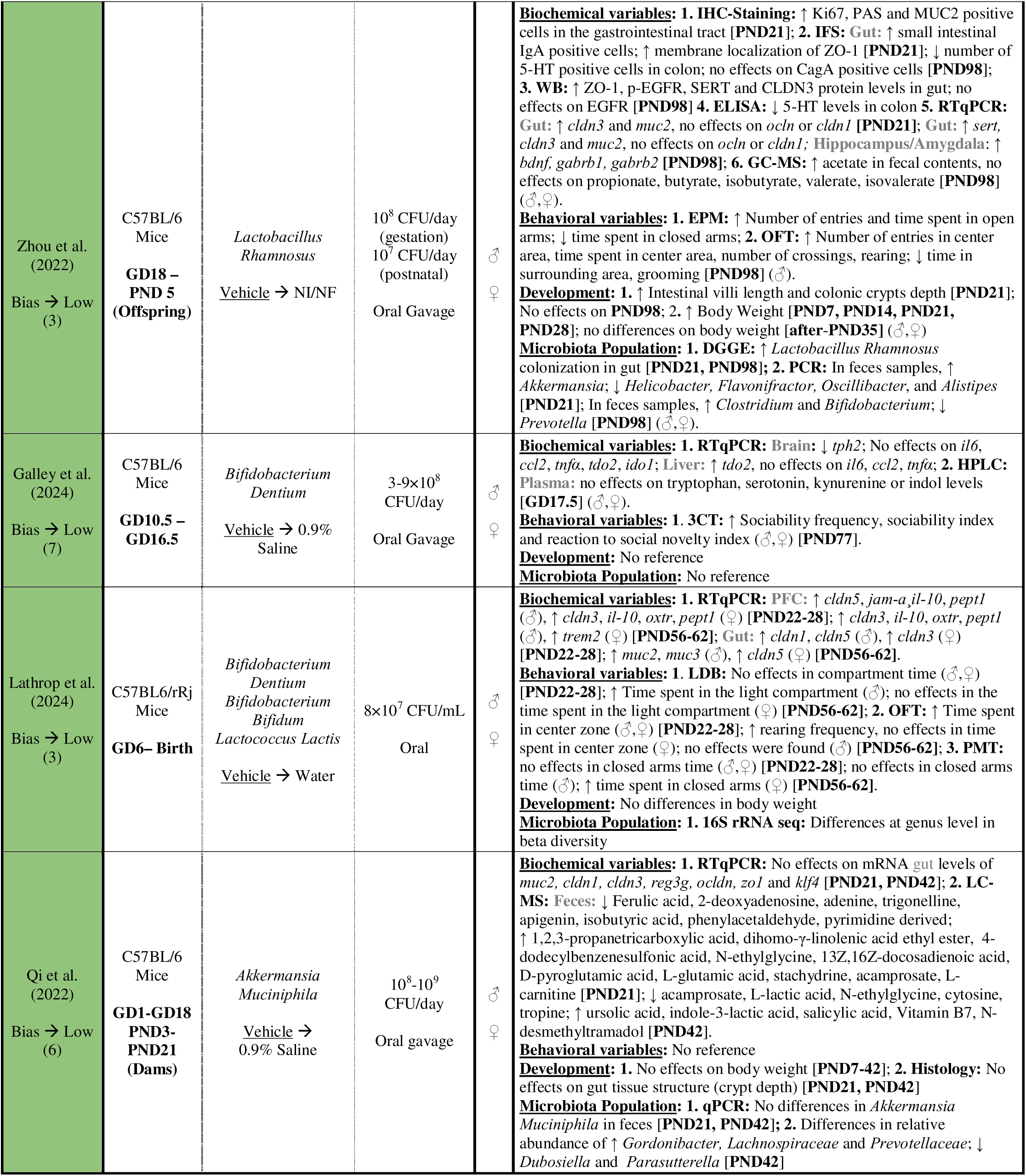
Summary of the materials, methods, and results found in the mouse studies included.

Abbr. PND: Postnatal day; GD: Gestational day; HPAC: High-Performance Liquid 228 Chromatography; GC-MS: Gas Chromatography Mass Spectrometry; RIA: Radioimmunoassay; IMA: Immunology Multiplex Assay; LC-MS:Liquid Chromatography-Mass Spectrometry; IFS: Immunofluorescence Staining; HPLC: High-Performance Liquid Chromatography; ELISA: Enzyme-Linked ImmunoSorbent Assay; EIA: Non-ELISA Enzyme Immunoassay; DGGE: Denaturing Gradient Gel Electrophoresis; WB: Western Blotting; IHC: Immunohistochemistry; 232 GCS: Golgi-Cox Staining; LDB: Light-Dark Box Test; FST: Forced Swim Test; MBT: Marble Burying Test; RSG: Repetitive Self-Grooming Test; MVF: Manual Von Frey Test; RTqPCR: Quantitative Reverse Transcription PCR; 3CT: Three-Chamber Test; EPM: Elevated Plus Maze; TST: Tail Suspension Test; OFT: Open Field Test; ROTAROD: RotaRod Performance Test; RT-PCRarray: Reverse Transcription PCR Array; PMT: Plus Maze Test; 16S rRNA seq: 16S Ribosomal RNA Sequencing.

### 3.2 General outcomes

All descriptive information regarding the methodology and results of each individual study is summarized in Table 1 (rat models) and Table 2 (mouse models). 5 of the 15 total studies (33.3%) used rat models while the rest of the 10 studies (66.7%) used mouse models. Furthermore, only 3 studies used only male sex, the rest of the studies used both sexes to test their supplementations. During the review, when sex has a significant impact on the results, the specific sex in which the observed effect occurs is indicated. Otherwise, it means that no differences were found, and the effect applies to both sexes. The systematic review revealed a wide variety of bacterial strains used in probiotic supplementation by the studies included, with *Lactobacillus* and *Bifidobacterium* genus being the most prominent. The most common strains include *L. Acidophilus, L. Fermentum*, *L. Rhamnosus*, *L. Plantarum, B. Lactis, B. Infantis, B. Longum, B. Breve* and *B. Bifidum* among others (see Table 1 and Table 2 for more detailed information), which were administered at various doses and routes of exposure (oral gavage and voluntary oral route by adding supplementation in drinking water). In addition, 9 of the studies included mixtures of multiple strains at the same time. Doses were measured in the studies in colony forming units (CFU) per day and per mL. The most common doses used in studies range from 10^6^ to 10^11^ CFU/day, however, some studies administered much larger doses of probiotic (e.g.: Barouei et al. (2012) used 3x10^9^ CFU/mL and 8 x 10^8^ CFU/mL in drinking water). The studies covered different times of administration, from early gestation to weaning. A total of 40% of the studies employed prenatal supplementation, while the remaining 60% combined both prenatal and postnatal treatments. The vehicle used was principally water, however, 2 studies used skimmed cow milk in order to facilitate absorption and other 2 studies used 0.9% saline. Lastly, only 3 studies had other excipients accompanied probiotic treatment. Wang et al. (2019) included Fructooligosaccharides (FOS) and maltodextrin, Kim and Lee (2022) 0.015 g/day of agarose and Azagra-Boronat et al. (2016) included maltodextrin.

### 3.3 Nervous system effects

The principal effects of prenatal bacterial strains administration during gestational period on central nervous system (CNS) and on peripheral tissue are classified and summarized in Table 3 and Table 4, respectively.

**Table 3.**
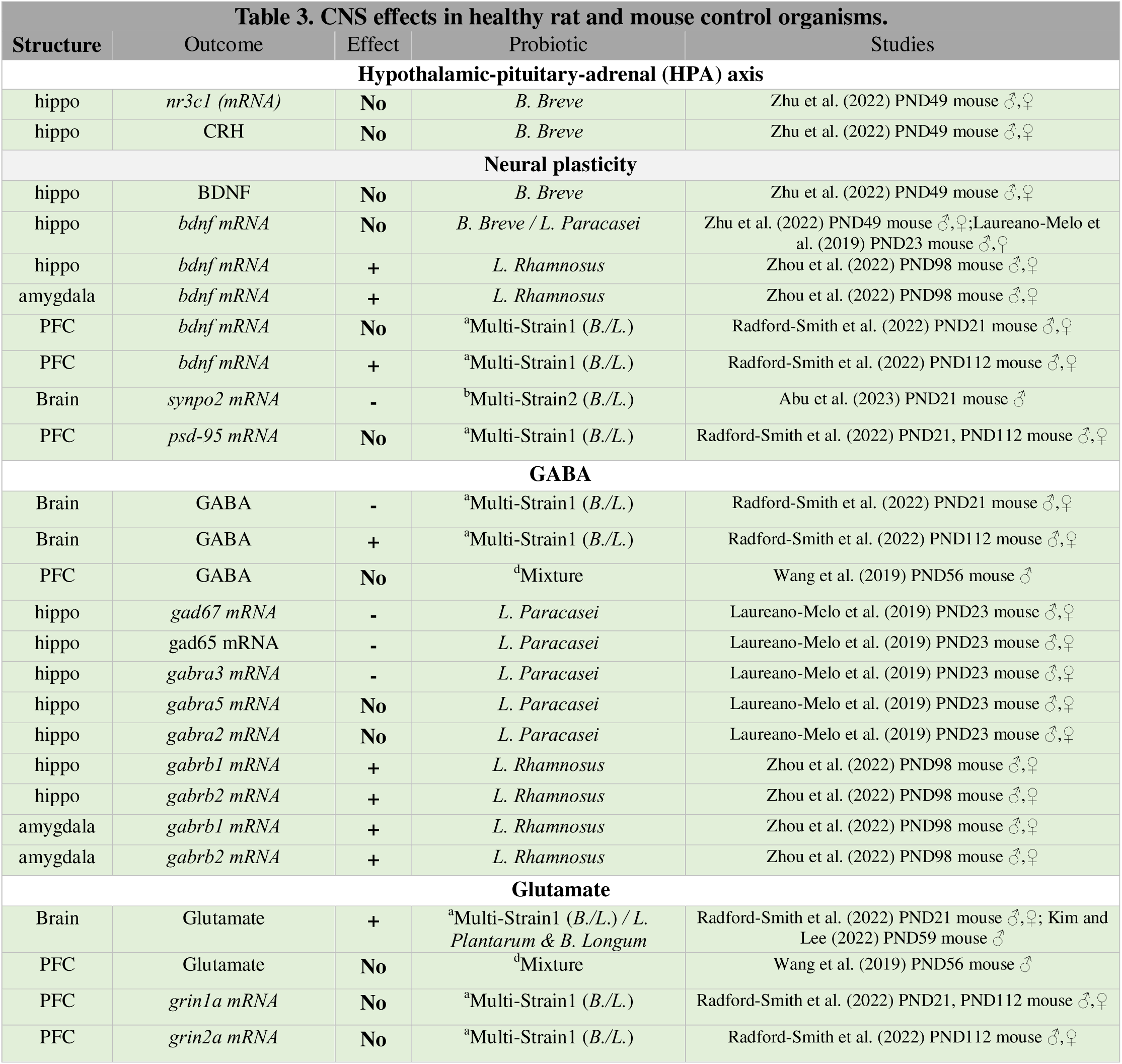
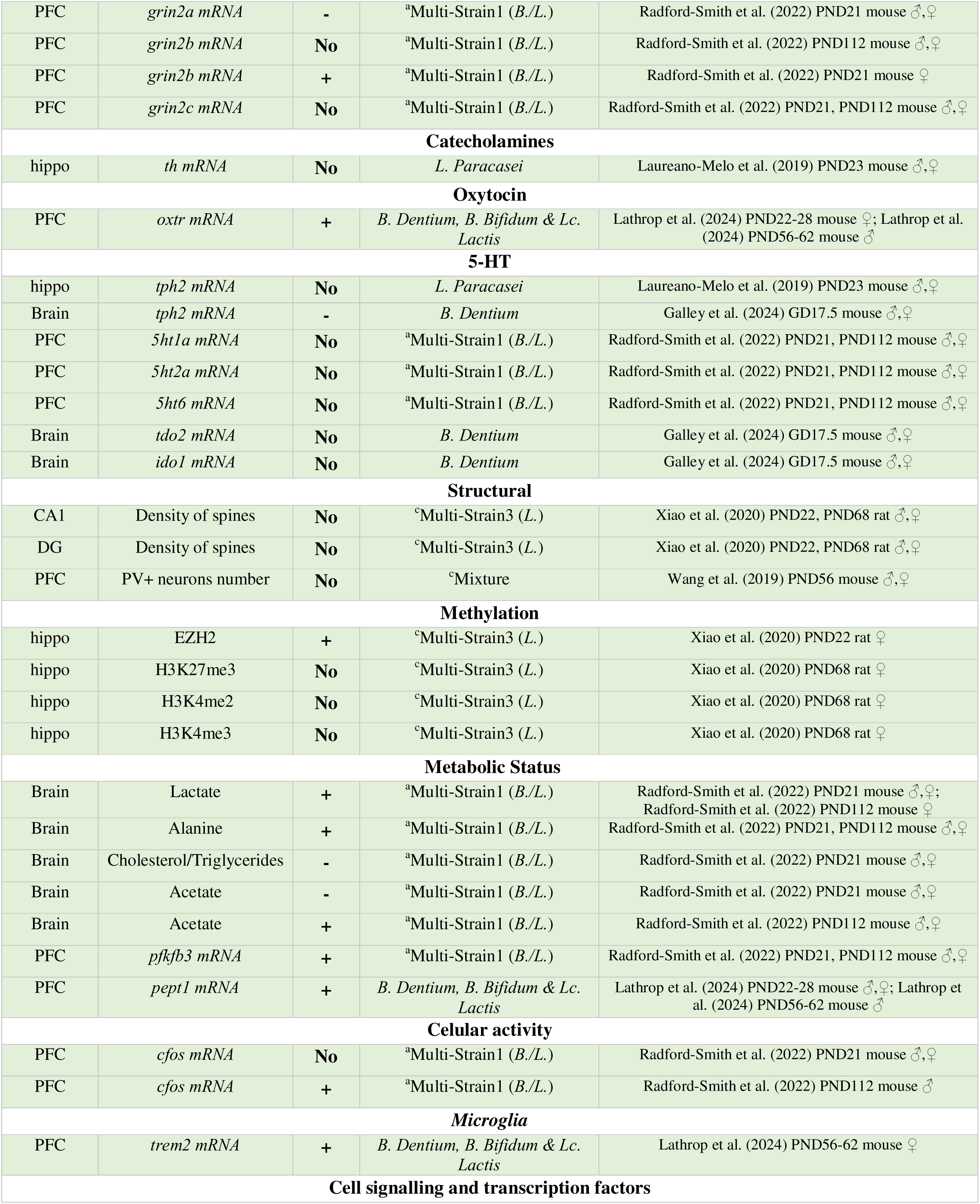
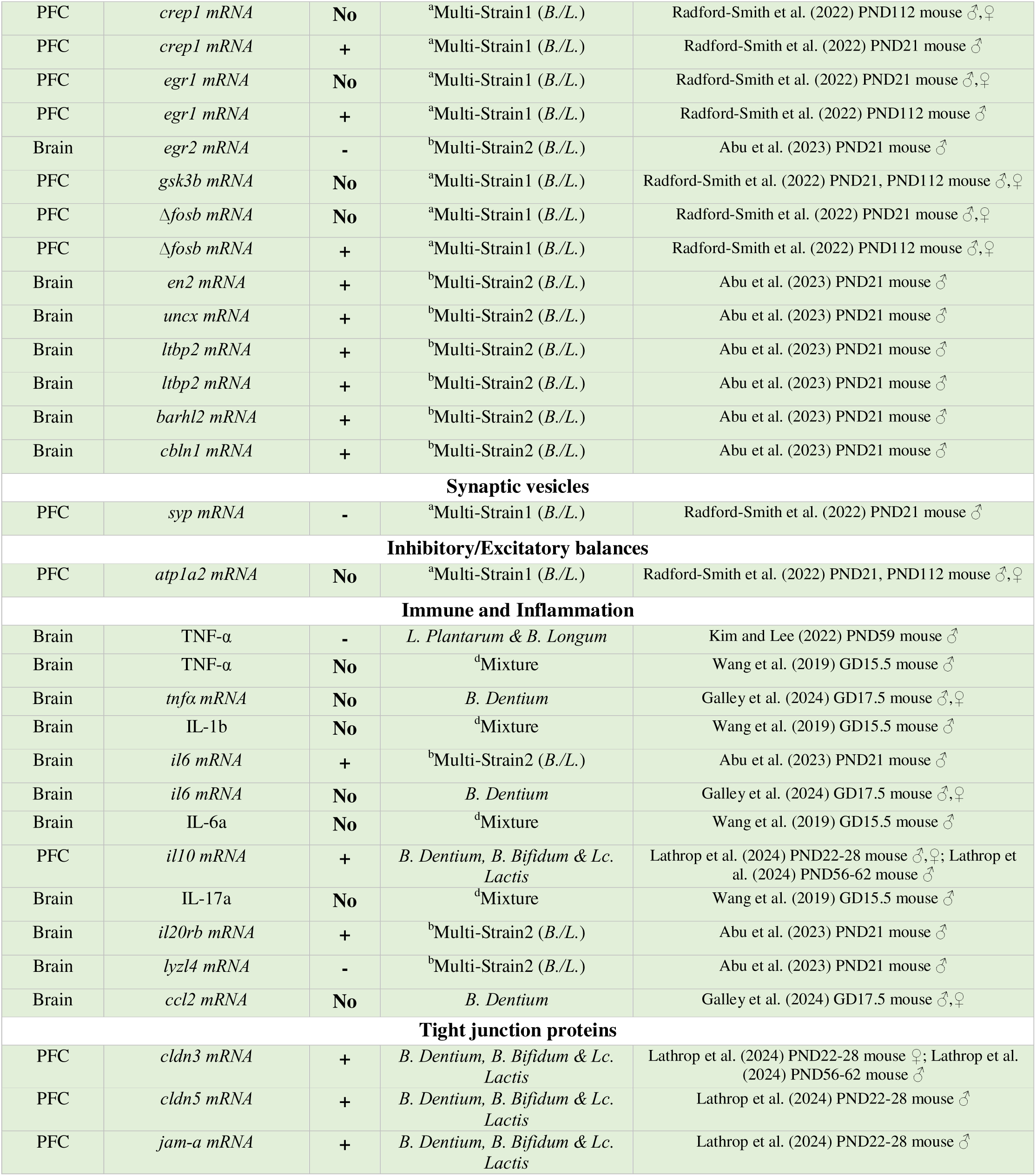
CNS effects in healthy rat and mouse control organisms.

Abbr. ^a^Multi-Strain1: B. Subtilis, B. Bifidum, B. Breve, B. Infantis, B. Longum, L. Acidophilus, L. Bulgaricus, L. Casei, L. Plantarum, L. Rhamnosus, L. Helveticus, L. Salivarius, Lc. Lactis and S. Thermophiles; bMulti-Strain2: L. Acidophilus, L. Bulgaricus, L. Paracasei, L. Plantarum, B. Breve, B. Infantis, B. Longum and S. Thermophilus; cMulti-Strain3: B. Longum, L. Acidophilus, L. Fermentum, L. Helveticus, L. Paracasei, L. Rhamnosus and S. Thermophilus; dMixture: 246 B. Bifidum, B. Infantis, L. Helveticus, FOS and Maltodextrin; PFC: Prefrontal Cortex; Hippo: Hippocampus; DG: Dental Gyrus.

**Table 4.** Peripheral effects in healthy rat and mouse control organisms.

| Structure | Outcome | Effect | Probiotic | Studies |
| --- | --- | --- | --- | --- |
| **Hypothalamic-pituitary-adrenal (HPA) axis** |  |  |  |  |
| Blood | Corticosterone | No | *L. Acidophilus*, *B. Lactis* & *L. Fermentum* / *B. Lactis* & *Propionibacterium Jensenii* / *B. Breve* | Hadizadeh et al. (2018) PND52 rat ♂; Barouei et al. (2012) PND86 rat ♂, ♀; Zhu et al. (2022) PND49 mouse ♀ |
| Blood | Corticosterone | + | *B. Lactis* & *Propionibacterium Jensenii* / *B. Breve* | Barouei et al. (2012) PND24 rat ♂, ♀; Zhu et al. (2022) PND49 mouse ♂ |
| Gut | *crhr1* mRNA | + | *B. Lactis* & *Propionibacterium Jensenii* | Moussavi et al. (2023) PND24 rat ♂, ♀ |
| Gut | *crhr1* mRNA | No | *B. Lactis* & *Propionibacterium Jensenii* | Moussavi et al. (2023) PND86 rat ♂, ♀ |
| Gut | *crhr2* mRNA | No | *B. Lactis* & *Propionibacterium Jensenii* | Moussavi et al. (2023) PND24, PND86 rat ♂, ♀ |
| **Achetylcholine** |  |  |  |  |
| Blood | Ach | No | *B. Breve* | Zhu et al. (2022) PND49 mouse ♂, ♀ |
| **Glutamate** |  |  |  |  |
| Feces | D-pyroglutamic acid | + | *A. Muciniphila* | Qi et al. (2022) PND21 mouse ♂, ♀ |
| Feces | L-glutamic acid | + | *A. Muciniphila* | Qi et al. (2022) PND21 mouse ♂, ♀ |
| Feces | Acamprosate | + | *A. Muciniphila* | Qi et al. (2022) PND21 mouse ♂, ♀ |
| Feces | Acamprosate | - | *A. Muciniphila* | Qi et al. (2022) PND42 mouse ♂, ♀ |
| **Gut mucus proteins** |  |  |  |  |
| Gut | *muc2* mRNA | + | *L. Rhamnosus* / *B. Dentium*, *B. Bifidum* & *Lc. Lactis* | Zhou et al. (2022) PND21, PND98 mouse ♂, ♀; Lathrop et al. (2024) PND56-62 mouse ♂ |
| Gut | *muc2* mRNA | No | *A. Muciniphila* | Qi et al. (2022) PND21, PND42 mouse ♂, ♀ |
| Gut | *muc3* mRNA | + | *B. Dentium*, *B. Bifidum* & *Lc. Lactis* | Lathrop et al. (2024) PND56-62 mouse ♂ |

| 5-HT |  |  |  |  |
| --- | --- | --- | --- | --- |
| Gut | *tph1 mRNA* | No | *B. Breve* | Zhu et al. (2022) PND49 mouse ♂,♀ |
| Gut | L-tryptophan | No | *B. Breve* | Zhu et al. (2022) PND49 mouse ♂,♀ |
| Blood | Tryptophan | No | *B. Dentium* | Galley et al. (2024) GD17.5 mouse ♂,♀ |
| Gut | 5-HIAA | No | *B. Breve* | Zhu et al. (2022) PND49 mouse ♂,♀ |
| Gut | 5-HTP | No | *B. Breve* | Zhu et al. (2022) PND49 mouse ♂,♀ |
| Gut | 5-HT | No | *B. Breve* | Zhu et al. (2022) PND49 mouse ♂,♀ |
| Blood | 5-HT | No | *B. Dentium* | Galley et al. (2024) GD17.5 mouse ♂,♀ |
| Gut | 5-HT | - | *L. Rhamnosus* | Zhou et al. (2022) PND98 mouse ♂,♀ |
| Gut | SERT | + | *L. Rhamnosus* | Zhou et al. (2022) PND98 mouse ♂,♀ |
| Gut | *sert mRNA* | + | *L. Rhamnosus* | Zhou et al. (2022) PND98 mouse ♂,♀ |
| Liver | *tdo2 mRNA* | + | *B. Dentium* | Galley et al. (2024) GD17.5 mouse ♂,♀ |
| Blood | Kynurenine | No | *B. Dentium* | Galley et al. (2024) GD17.5 mouse ♂,♀ |
| Cell signalling and transcription factors |  |  |  |  |
| Gut | p-EGFR | + | *L. Rhamnosus* | Zhou et al. (2022) PND98 mouse ♂,♀ |
| Gut | *klf4 mRNA* | No | *A. Muciniphila* | Qi et al. (2022) PND21, PND42 mouse ♂,♀ |
| Gut | *cfos mRNA* | No | *B. Breve* | Zhu et al. (2022) PND49 mouse ♂,♀ |
| Immune and Inflammation |  |  |  |  |
| Feces | IgA | + | *L. Fermentum* | Azagra-Boronat et al. (2016) PND14 rat ♂,♀ |
| Blood | IgA | - | *B. Lactis* & *P. Jensenii* | Barouei et al. (2012) PND24 rat ♂,♀ |
| Blood | IgA | No | *B. Lactis* & *P. Jensenii* | Barouei et al. (2012) PND86 rat ♂,♀ |
| Blood | IgM | No | *L. Fermentum* | Azagra-Boronat et al. (2016) PND14 rat ♂,♀ |
| Feces | IgM | No | *L. Fermentum* | Azagra-Boronat et al. (2016) PND14 rat ♂,♀ |
| Feces | IgG | No | *L. Fermentum* | Azagra-Boronat et al. (2016) PND14 rat ♂,♀ |
| Blood | IgG1 | No | *L. Fermentum* | Azagra-Boronat et al. (2016) PND14 rat ♂,♀ |
| Blood | IgG2a | + | *L. Fermentum* | Azagra-Boronat et al. (2016) PND14 rat ♂,♀ |
| Blood | IgG2b | No | *L. Fermentum* | Azagra-Boronat et al. (2016) PND14 rat ♂,♀ |
| Blood | IgG2c | - | *L. Fermentum* | Azagra-Boronat et al. (2016) PND14 rat ♂,♀ |
| Blood | G-CSF | + | °Multi-Strain3 (*L.*) | Xiao et al. (2020) PND68 rat ♀ |
| Blood | IL-4 | No | *L. Fermentum* | Azagra-Boronat et al. (2016) PND14 rat ♂,♀ |
| Blood | IL-6 | No | °Multi-Strain3 (*L.*) / *L. Fermentum* | Xiao et al. (2020) PND68 rat ♀; Azagra-Boronat et al. (2016) PND14 rat ♂,♀ |
| Liver | *il6 mRNA* | No | *B. Dentium* | Galley et al. (2024) GD17.5 mouse ♂,♀ |
| Blood | IL-10 | No | °Multi-Strain3 (*L.*) / *L. Fermentum* | Xiao et al. (2020) PND68 rat ♀; Azagra-Boronat et al. (2016) PND14 rat ♂,♀ |
| Blood | IL-12 | No | *L. Fermentum* | Azagra-Boronat et al. (2016) PND14 rat ♂,♀ |
| Blood | TNF-α | No | °Multi-Strain3 (*L.*) / *L. Fermentum* | Xiao et al. (2020) PND68 rat ♀; Azagra-Boronat et al. (2016) PND14 rat ♂,♀ |
| Liver | *tnfa mRNA* | No | *B. Dentium* | Galley et al. (2024) GD17.5 mouse ♂,♀ |
| Blood | IFN-γ | No | °Multi-Strain3 (*L.*) / *L. Fermentum* | Xiao et al. (2020) PND68 rat ♀; Azagra-Boronat et al. (2016) PND14 rat ♂,♀ |
| Liver | *ccl2 mRNA* | No | *B. Dentium* | Galley et al. (2024) GD17.5 mouse ♂,♀ |
| Gut | *reg3g mRNA* | No | *A. Muciniphila* | Qi et al. (2022) PND21, PND42 mouse ♂,♀ |
| Feces | Ursolic acid | + | *A. Muciniphila* | Qi et al. (2022) PND42 mouse ♂,♀ |
| Enteric nervous system |  |  |  |  |
| Gut | *ngf mRNA* | No | *B. Lactis* & *P. Jensenii* | Moussavi et al. (2023) PND24, PND86 rat ♂,♀ |

| Gut | Substance P | No | *B. Breve* | Zhu et al. (2022) PND49 mouse ♂,♀ |
| --- | --- | --- | --- | --- |
| **Fatty acids** |  |  |  |  |
| Blood | Palmitic acid | - | *L. Fermentum* | Azagra-Boronat et al. (2016) PND14 rat ♂,♀ |
| Blood | Total saturated fatty acids | - | *L. Fermentum* | Azagra-Boronat et al. (2016) PND14 rat ♂,♀ |
| Blood | Linoleic acid | + | *L. Fermentum* | Azagra-Boronat et al. (2016) PND14 rat ♂,♀ |
| Blood | Eicosadinoic acid | + | *L. Fermentum* | Azagra-Boronat et al. (2016) PND14 rat ♂,♀ |
| Blood | Margaric fatty acids | + | *L. Fermentum* | Azagra-Boronat et al. (2016) PND14 rat ♂,♀ |
| Blood | Behenic fatty acids | + | *L. Fermentum* | Azagra-Boronat et al. (2016) PND14 rat ♂,♀ |
| Feces | Isobutyric acid | - | *A. Muciniphila* | Qi et al. (2022) PND21 mouse ♂,♀ |
| Feces | dihomo-γ-linolenic acid ethyl ester | + | *A. Muciniphila* | Qi et al. (2022) PND21 mouse ♂,♀ |
| **SCFAs** |  |  |  |  |
| Feces | Butyrate | + | <sup>a</sup>Multi-Strain1 (*B./L.*) | Radford-Smith et al. (2022) PND21, PND112 mouse ♂,♀ |
| Feces | Butyrate | **No** | *L. Rhamnosus* | Zhou et al. (2022) PND98 mouse ♂,♀ |
| Feces | Propionate | + | <sup>a</sup>Multi-Strain1 (*B./L.*) | Radford-Smith et al. (2022) PND21, PND112 mouse ♂,♀ |
| Feces | Propionate | **No** | *L. Rhamnosus* | Zhou et al. (2022) PND98 mouse ♂,♀ |
| Feces | Acetate | + | *L. Rhamnosus* | Zhou et al. (2022) PND98 mouse ♂,♀ |
| Blood | Acetate | - | <sup>a</sup>Multi-Strain1 (*B./L.*) | Radford-Smith et al. (2022) PND21 mouse ♂,♀;<br>Radford-Smith et al. (2022) PND112 mouse ♂ |
| Liver | Acetate | + | <sup>a</sup>Multi-Strain1 (*B./L.*) | Radford-Smith et al. (2022) PND21 mouse ♂,♀ |
| Feces | Isobutyrate | **No** | *L. Rhamnosus* | Zhou et al. (2022) PND98 mouse ♂,♀ |
| Feces | Valerate | **No** | *L. Rhamnosus* | Zhou et al. (2022) PND98 mouse ♂,♀ |
| Feces | Isovalerate | **No** | *L. Rhamnosus* | Zhou et al. (2022) PND98 mouse ♂,♀ |
| Feces | Total SCFAs | **No** | *B. Breve* | Zhu et al. (2022) PND49 mouse ♂,♀ |
| **Metabolic Status** |  |  |  |  |
| Blood | Lactate | + | <sup>a</sup>Multi-Strain1 (*B./L.*) | Radford-Smith et al. (2022) PND21 mouse ♂,♀ |
| Liver | Lactate | + | <sup>a</sup>Multi-Strain1 (*B./L.*) | Radford-Smith et al. (2022) PND21 mouse ♂,♀;<br>Radford-Smith et al. (2022) PND112 mouse ♀ |
| Blood | Alanine | + | <sup>a</sup>Multi-Strain1 (*B./L.*) | Radford-Smith et al. (2022) PND21 mouse ♂,♀ |
| Blood | Alanine | - | <sup>a</sup>Multi-Strain1 (*B./L.*) | Radford-Smith et al. (2022) PND112 mouse ♂,♀ |
| Liver | Alanine | + | <sup>a</sup>Multi-Strain1 (*B./L.*) | Radford-Smith et al. (2022) PND112 mouse ♂,♀ |
| Feces | Alanine | - | <sup>a</sup>Multi-Strain1 (*B./L.*) | Radford-Smith et al. (2022) PND112 mouse ♂ |
| Blood | Glucose | - | <sup>a</sup>Multi-Strain1 (*B./L.*) | Radford-Smith et al. (2022) PND21 mouse ♂,♀;<br>Radford-Smith et al. (2022) PND112 mouse ♂ |
| Liver | Glucose | + | <sup>a</sup>Multi-Strain1 (*B./L.*) | Radford-Smith et al. (2022) PND112 mouse ♂ |
| Liver | Taurine | - | <sup>a</sup>Multi-Strain1 (*B./L.*) | Radford-Smith et al. (2022) PND21 mouse ♂,♀ |
| Feces | Ethanol | - | <sup>a</sup>Multi-Strain1 (*B./L.*) | Radford-Smith et al. (2022) PND21 mouse ♂,♀ |
| Feces | Ethanol | + | <sup>a</sup>Multi-Strain1 (*B./L.*) | Radford-Smith et al. (2022) PND112 mouse ♂ |
| Feces | Alanine | - | <sup>a</sup>Multi-Strain1 (*B./L.*) | Radford-Smith et al. (2022) PND21 mouse ♂,♀ |
| Blood | VLDL/TG | - | <sup>a</sup>Multi-Strain1 (*B./L.*) | Radford-Smith et al. (2022) PND112 mouse ♂ |
| Liver | Creatine | - | <sup>a</sup>Multi-Strain1 (*B./L.*) | Radford-Smith et al. (2022) PND112 mouse ♀ |
| Liver | Taurine | + | <sup>a</sup>Multi-Strain1 (*B./L.*) | Radford-Smith et al. (2022) PND112 mouse ♂,♀ |
| Blood | Indol | **No** | *B. Dentium* | Galley et al. (2024) GD17.5 mouse ♂,♀ |
| Feces | 2-deoxyadenosine | - | *A. Muciniphila* | Qi et al. (2022) PND21 mouse ♂,♀ |
| Feces | Adenosine | - | *A. Muciniphila* | Qi et al. (2022) PND21 mouse ♂,♀ |
| Feces | Phenylacetaldehyde | - | *A. Muciniphila* | Qi et al. (2022) PND21 mouse ♂,♀ |
| Feces | Pyrimidine derived | - | *A. Muciniphila* | Qi et al. (2022) PND21 mouse ♂,♀ |
| Feces | 1,2,3-propanetricarboxylic acid | + | *A. Muciniphila* | Qi et al. (2022) PND21 mouse ♂,♀ |
| Feces | N-ethylglycine | + | *A. Muciniphila* | Qi et al. (2022) PND21 mouse ♂,♀ |
| Feces | L-carnitine | + | *A. Muciniphila* | Qi et al. (2022) PND21 mouse ♂,♀ |
| Feces | L-lactic acid | - | *A. Muciniphila* | Qi et al. (2022) PND42 mouse ♂,♀ |
| Feces | N-ethylglycine | - | *A. Muciniphila* | Qi et al. (2022) PND42 mouse ♂,♀ |
| Feces | Cytosine | - | *A. Muciniphila* | Qi et al. (2022) PND42 mouse ♂,♀ |
| Feces | Tropine | - | *A. Muciniphila* | Qi et al. (2022) PND42 mouse ♂,♀ |
| Feces | Vitamin B7 | + | *A. Muciniphila* | Qi et al. (2022) PND42 mouse ♂,♀ |
| Feces | Indole-3-lactic acid | + | *A. Muciniphila* | Qi et al. (2022) PND42 mouse ♂,♀ |
| Feces | Salicylic acid | + | *A. Muciniphila* | Qi et al. (2022) PND42 mouse ♂,♀ |
| Feces | N-desmethyltramadol | + | *A. Muciniphila* | Qi et al. (2022) PND42 mouse ♂,♀ |
| Feces | Trigoneline | - | *A. Muciniphila* | Qi et al. (2022) PND21 mouse ♂,♀ |
| **Oxidative stress** |  |  |  |  |
| Liver | GSSG | + | <sup>a</sup>Multi-Strain1 (*B./L.*) | Radford-Smith et al. (2022) PND21 mouse ♂,♀ |
| Feces | Ferulic acid | - | *A. Muciniphila* | Qi et al. (2022) PND21 mouse ♂,♀ |
| Feces | Apigenin | - | *A. Muciniphila* | Qi et al. (2022) PND21 mouse ♂,♀ |
| **Gut structure and function** |  |  |  |  |
| Gut | Epithelial damage and inflammation | **No** | <sup>b</sup>Multi-Strain2 (*B./L.*) | Abu et al. (2023) PND21 mouse ♂ |
| Gut | Ki67 positive cells | + | *L. Rhamnosus* | Zhou et al. (2022) PND21 mouse ♂,♀ |
| Gut | PAS positive cells | + | *L. Rhamnosus* | Zhou et al. (2022) PND21 mouse ♂,♀ |
| Gut | MUC2 positive cells | + | *L. Rhamnosus* | Zhou et al. (2022) PND21 mouse ♂,♀ |
| Gut | Small IgA positive cells | + | *L. Rhamnosus* | Zhou et al. (2022) PND21 mouse ♂,♀ |
| Gut | Membrane location of ZO-1 | + | *L. Rhamnosus* | Zhou et al. (2022) PND21 mouse ♂,♀ |
| Gut | 5-HT positive cells | + | *L. Rhamnosus* | Zhou et al. (2022) PND98 mouse ♂,♀ |
| Gut | CagA positive cells | **No** | *L. Rhamnosus* | Zhou et al. (2022) PND98 mouse ♂,♀ |
| Gut | Villi length | + | *L. Rhamnosus* | Zhou et al. (2022) PND21 mouse ♂,♀ |
| Gut | Villi length | **No** | *L. Rhamnosus* | Zhou et al. (2022) PND98 mouse ♂,♀ |
| Gut | Crypts depth | + | *L. Rhamnosus* | Zhou et al. (2022) PND21 mouse ♂,♀ |
| Gut | Crypts depth | **No** | *L. Rhamnosus* / *A. Muciniphila* | Zhou et al. (2022) PND98 mouse ♂,♀; Qi et al. (2022) PND21, PND42 mouse ♂,♀ |
| **Tight junction proteins** |  |  |  |  |
| Gut | ZO-1 | + | *L. Rhamnosus* | Zhou et al. (2022) PND98 mouse ♂,♀ |
| Gut | *zo1* mRNA | **No** | *A. Muciniphila* | Qi et al. (2022) PND21, PND42 mouse ♂,♀ |
| Gut | CLDN3 | + | *L. Rhamnosus* | Zhou et al. (2022) PND98 mouse ♂,♀ |
| Gut | *cldn3* mRNA | + | *L. Rhamnosus* / *B. Dentium*, *B. Bifidum* & *Lc. Lactis* | Zhou et al. (2022) PND21, PND98 mouse ♂,♀; Lathrop et al. (2024) PND22-28 mouse ♀ |
| Gut | *cldn3* mRNA | **No** | *A. Muciniphila* | Qi et al. (2022) PND21, PND42 mouse ♂,♀ |
| Gut | *ocldn* mRNA | **No** | *L. Rhamnosus* / *A. Muciniphila* | Zhou et al. (2022) PND21, PND98 mouse ♂,♀; Qi et al. (2022) PND21, PND42 mouse ♂,♀ |
| Gut | *cldn1* mRNA | **No** | *L. Rhamnosus* / *B. Dentium*, *B. Bifidum* & *Lc. Lactis* / *A.* | Zhou et al. (2022) PND21, PND98 mouse ♂,♀; Lathrop et al. (2024) PND22-28 mouse ♀; Qi et al. (2022) PND21, PND42 mouse ♂,♀ |
|  |  |  | *Muciniphila* |  |
| Gut | *cldn1* mRNA | + | *B. Dentium*, *B. Bifidum* & *Lc. Lactis* | Lathrop et al. (2024) PND22-28 mouse ♂ |
| Gut | *cldn5* mRNA | + | *B. Dentium*, *B. Bifidum* & *Lc. Lactis* | Lathrop et al. (2024) PND22-28 mouse ♂; Lathrop et al. (2024) PND56-62 mouse ♂ |

Abbr. ^a^Multi-Strain1: B. Subtilis, B. Bifidum, B. Breve, B. Infantis, B. Longum, L. Acidophilus, L.
Bulgaricus, L. Casei, L. Plantarum, L. Rhamnosus, L. Helveticus, L. Salivarius, Lc. Lactis and S.
Thermophiles; bMulti-Strain2: L. Acidophilus, L. Bulgaricus, L. Paracasei, L. Plantarum, B. Breve, B.
Infantis, B. Longum and S. Thermophilus; cMulti-Strain3: B. Longum, L. Acidophilus, L. Fermentum, L.
Helveticus, L. Paracasei, L. Rhamnosus and S. Thermophilus; dMixture: B. Bifidum, B. Infantis, L.
Helveticus, FOS and Maltodextrin.

#### 3.3.1 Hypothalamic-Pituitary-Adrenal axis

Only 4 of the 15 studies (26.7%) reported findings about the Hypothalamic-Pituitary- Adrenal (HPA) axis. Zhu et al. (2022) found that a full-gestation supplementation with *B. Breve* did not alter either the gene expression of *nr3c1* (glucocorticoid receptor) or the levels of the corticotropin-releasing hormone (CRH) in the hippocampus of PND49 mice. One study analysed *crh* receptors mRNA levels in gut tissue and discovered that while *crhr1* was up-regulated in PND24 rats, it was not affected in PND86 rats exposed pre- and postnatally to *B. Lactis* and *P. Jensenii*. Neither were the mRNA levels of *crhr2* affected at any age (Moussavi et al., 2023). Finally, 2 of the 3 studies that analysed corticosterone (CORT) blood levels found an increase both in PND24 rats (Barouei et al., 2012) and PND49 male mice (Zhu et al., 2022). In contrast, no effects in CORT blood levels were found in those same studies when the rats reached adulthood, nor in female mice. Also, Hadizadeh et al. (2018) did not find effects in PND52 male rats treated until GD14 with *L. Acidophilus*, *L. Fermentum* and *B. Lactis*.

#### 3.3.2 Neural plasticity

A total of 4 of 15 studies (26.7%) showed results about brain-derived neurotrophic factor (BDNF) function. The only one that examined BDNF at the protein level did not found alterations in hippocampus of PND49 mice after prenatal *B. Breve* (Zhu et al., 2022). However, the results regarding their mRNA levels are controversial. While Zhu et al. (2022) and Laureano-Melo et al. (2019) did not find *bdnf* alterations with *L. Paracasei* in mice’s hippocampus, an increase in *bdnf* mRNA levels was found in hippocampus and amygdala in PND98 mice supplemented with *L. Rhamnosus*. Lastly, in PFC no effects were obtained when a multi-strain probiotic (principally composed of *Lactobacillus* and *Bifidobacterium* genus) was administered from GD0.5 until weaning in mice at PND21, but an up-regulation of BDNF was noted at PND112. Only one study analysed the expression of *synpo2* (synaptopodin 2) finding a down-regulation in whole brain tissue. Lastly, methylation-related molecules were observed at PND22 in the hippocampus of female rats primarily supplemented with *Lactobacillus* strains from conception until weaning, with an increase in EZH2 levels. However, no significant effects were noted for H3K27me3, H3K4me2, and H3K4me3 at PND68 (Xiao et al., 2020).

#### 3.3.3 GABAergic system

26.7% studies analysed how prenatal probiotic treatments impact in mice central GABAergic system formation. Of those 4 studies, 2 examine GABA levels in CNS. In mice pre- and postnatally supplemented with multi-strain probiotic, a decrease of GABA in the whole brain was observed at PND21. However, the opposite effect was found in PND112, an increase in GABA levels (Radford-Smith et al., 2022). On the other hand, Wang et al. (2019) did not find GABA levels alterations in PFC of PND56 male mice after gestational treatment with *B. Bifidum*, *B. Infantis*, *L. Helveticus*, FOS and maltodextrin. Finally, the other 2 studies analysed the gene expression of glutamic acid decarboxylase (GAD) and GABA-A receptor subunits. Prenatal *L. Paracasei* led to a down-regulation of hippocampal *gad67*, *gad65* and *gabra3* in PND23 mice, while no effects were observed on *gabra2* and *gabra5* (Laureano-Melo et al., 2019). In contrast, *L. Rhamnosus* supplementation (between GD18 and PND5) caused an up-regulation of *gabrb1* and *gabrb2* in both hippocampus and amygdala of PND98 mice.

#### 3.3.4 Glutamatergic system

Glutamate system related molecules were assessed in 20% of the studies. While glutamate was increased in overall brain of PND21 mice treated with multi-strain probiotic (Radford-Smith et al., 2022) and PND59 male mice prenatally supplemented with *L. Plantarum* and *B. Longum* (Kim and Lee, 2022), no effects were found in PFC of PND56 male mice prenatally treated with *B. Bifidum*, *B. Infantis*, *L. Helveticus*, FOS and maltodextrin (Wang et al., 2019). In this sense, Radford-Smith et al. (2022) also analysed the impact in PFC of a pre- and postnatal multi-strain probiotic (primarily *Lactobacillus* and *Bifidobacterium*). In PND21 mice, there was an up-regulation of *grin2b* (only in males) and a down-regulation of *grin2a*; an effect that disappears in PND112. No effects were found in *grin1a* or *grin2c* at any age. Furthermore, Qi et al. (2022) revealed several notable changes in glutamate-related metabolites in fecal samples of mice following a pre- and postnatal treatment with *Akkermansia Muciniphila*. In PND21, D-pyroglutamic acid, acamprosate and L-glutamic acid showed a marked increase. However, by PND42, a decrease in acamprosate levels was recorded in the same group of mice.

#### 3.3.5 Serotonergic system

A total of 33.3% of the studies explored the function of serotonergic system in mice model. For the mRNA levels of tryptophan hydroxylase 2 gene (*tph2*), varied results were observed. Pre- and postnatal supplementation with *L. Paracasei* did not affects its expression in hippocampus of PND23 mice (Laureano-Melo., 2019). However, in the brains of prenatal mice (GD17.5), a brief treatment with *B. dentium* between GD10.5 and GD16.5 resulted in the downregulation of *tph2*. This treatment did not affect the expression of *ido1* and *tdo2* in the brain or the levels of kynurenine, tryptophan or serotonin (5-HT) in blood; however, an increase in *tdo2* mRNA was observed in the liver (Galley et al., 2024). Radford-Smith et al. (2022) observed that pre- and postnatal multi-strain probiotic did not affect gene expression of *5ht1a*, *5ht2a* and *5ht6* in PFC of young and adult mice. In the gut, no significant effects were observed for *tph1* mRNA, L-tryptophan, 5-HIAA, 5-HTP, and 5-HT in PND49 mice prenatally treated with *B. Breve* (Zhu et al., 2022). Conversely, a decrease in 5-HT and increases in SERT and *sert* mRNA levels were noted at PND98 after *L. Rhamnosus* supplementation (Zhou et al., 2022).

#### 3.3.6 Other neurotransmitter systems and central nervous effects

At the structural level, there were no significant effects on the density of spines in the CA1 and dentate gyrus (DG) regions of the hippocampus at PND22 and PND68 rats (Xiao et al., 2020). Additionally, there was no observed effect on the number of parvalbumin-positive (PV+) neurons in the PFC at PND56 in mice (Wang et al., 2019). Regarding cell signalling and transcription factors in the context of prenatal and postnatal exposure to a multi-strain probiotic containing various *Lactobacillus* and *Bifidobacterium* species until weaning, in the PFC, *crep1* mRNA levels showed no effect at PND112 mice, but an increase was noted at PND21 in male mice (Radford-Smith et al., 2022). The *egr1* mRNA levels were unchanged at PND21, while an increase was observed at PND112 in male mice (Radford-Smith et al., 2022). In contrast, *egr2* decreased in the brain at PND21 (Abu et al., 2023). Other markers in the PFC, such as *gsk3b* and Δ*fosb*, showed no significant effects at PND21 and PND112, except for an increase in Δ*fosb* and *atp1a2* at PND112 (Radford-Smith et al., 2022).

Additionally, various mRNAs, including *en2*, *uncx*, *ltbp2*, *barhl2*, and *cbln1* demonstrated increases in the overall brain of mice treated with multi-strain probiotic at PND21 (Abu et al., 2023).

Although the studies did not analyse catecholamines, Laurenano-Melo et al. (2019) did not found effects in tyrosine hydroxylase (*th*) expression in hippocampus after *L. Paracasei* pre- and postnatal supplementation in PND23 mice. Regarding the oxytocinergic system, Lathrop et al. (2024) observed that prenatal supplementation with *B. Dentium, B. Bifidum, and L. Lactis* produced an up-regulation of oxytocin receptors (oxtr) in the mice PFC, with differential effects between males and females. In males, the upregulation occurred between PND56-62, while in females, it was observed between PND22-28. Lastly, other effects in *syp* (down-regulation) and *trem2* (up- regulation) were observed at specific time points (see Table 2). Acetylcholine blood levels were unchanged in PND49 mice prenatal supplemented with *B. Breve* (Zhu et al., 2022).

### 3.4 Inflammation, immune system and oxidative stress

The systematic review revealed significant findings related to immune and inflammatory markers. 60% of the studies included in the review analysed the immune system, with 5 examining these markers in the brain and 5 in peripheral tissues. While *L. Plantarum y B. Longum* gestational supplementation decreased TNF-α levels in overall brain of PND59 male mice (Kim and Lee, 2022), no effect was observed at GD15.5 or in *tnfa* expression at GD17.5 (Wang et al., 2019; Galley et al., 2024). No changes were noted for IL-1b, IL-6a, or IL-17a at GD15.5 (Wang et al., 2019), though *il6* mRNA increased at PND21 in male mice multi-strain supplemented (Abu et al., 2023) and remained unchanged at GD17.5 (Galley et al., 2024). After prenatal *B. Dentium, B. Bifidum* and *L. Lactis*, *il10* mRNA showed an increase at PND22-28 and PND56-62 in mice PFC (Lathrop et al., 2024). Furthermore, *il20rb* mRNA increased, while *lyzl4* decreased at PND21 (Abu et al., 2023). Lastly, *ccl2* mRNA showed no effect at GD17.5 (Galley et al., 2024), while *jam-a* mRNA increased in the PFC at PND22-28 (Lathrop et al., 2024).

Regarding to peripheral tissues, *L. Fermentum* pre- and postnatal treatment in Wistar Rats increased IgA levels in feces and IgG2a in blood and decreased IgG2c blood levels at PND14. No effects were found on blood levels of IgM, IgG1 and IgG2b or feces levels of IgM or IgG (Azagra-Boronat et al., 2016). In contrast, *B. Lactis* and *P. Jensenii* treatment decreased IgA rat blood levels at PND24, effects that were lost at PND86 (Barouei et al., 2012). Furthermore, *Lactobacillus* multi-strain gestational and early postnatal probiotic increase G-CSF levels in PND68 rats (Xiao et al., 2020).

Azagra-Boronat et al. (2016), which utilized a pre- and postnatal treatment in rats with *L. Fermentum*, found no significant effects on blood levels of IL-4, IL-6, IL-10, IL-12, TNF-α, and IFN-γ in both male and female rats at PND14. Similarly, Xiao et al. (2020) observed no effects on blood IL-6, IL-10, TNF-α, and IFN-γ levels in female rats at PND68 when applying a multi-strain probiotic treatment. In the liver, Galley et al. (2024) reported no effects on *il6* and *tnfa* mRNA levels in male and female mice at GD17.5 after prenatal treatment with *B. Dentium*. Qi et al. (2022) investigated a pre- and postnatal supplementation with *A. Municiphila* in mice and found no effect on gut *reg3g* mRNA levels but reported an increase in fecal ursolic acid levels in PND42 mice.

Regarding the influence of probiotics on oxidative stress, not many results have been found. Radford-Smith et al. (2022) reported an increase in liver GSSG levels in mice at PND21 exposed to *Lactobacillus* and *Bifidobacterium* multi-strain probiotic. On the other hand, Qi et al. (2022) found that *A. Muciniphila* decrease in fecal ferulic acid and apigenin levels in mice at the same age.

### 3.5 Fatty acids and SCFAs

A 33.3% of the studies showed results about changes in fatty acid concentration, of which 3 studies are specific to SCFAs. In the study conducted by Azagra-Boronat et al. (2016) on PND14 rats, *L. Fermentum* pre- and postnatal supplementation generated a decrease in palmitic acid and total saturated fatty acids in blood samples, while levels of linoleic acid, eicosadinoic acid, margaric fatty acids, and behenic fatty acids increased. Similarly, Qi et al. (2022) conducted research on PND21 mice, where a decrease in isobutyric acid was reported in feces, along with an increase in dihomo-γ-linolenic acid ethyl ester. Concerning SCFAs analysis, controversial results were obtained about the influence of gestational probiotics in its levels in peripheral tissues. While Radford- Smith et al. (2022) found an increase in butyrate and propionate feces levels in both PND21 and PND112 mice exposed during gestation and lactation to multi-strain probiotic, Zhou et al. (2022) did not find effects in these variables when PND98 mouse *L. Rhamnosus*-exposed is evaluated, however, acetate levels were increased in feces in this latter study. Radford-Smith et al. (2022), for his part, finds that the levels of acetate decrease in blood samples of PND21 for both sexes and PND112 for male mice, and increase in the liver of PND21 mice. On the other hand, *L. Rhamnosus* did not alter the fece levels of isobutyrate, valerate or isovalerate in PND98 mice (Zhou et al., 2022) and *B. Breve* gestational treatment did not change the total SCFAs concentration in feces of PND49 mice (Zhu et al., 2022).

### 3.6 Metabolic status

Of the studies that analysed metabolic outcomes (26.7% of total sample), only 2 are in brain tissue. In young mice (PND21), multi-strain probiotic induced an increase in lactate and alanine levels and a decrease in cholesterol/triglycerides ratio and acetate levels in overall brain. In adult mice (PND112) this probiotic also increased lactate (only in females) and alanine levels, but in this case acetate levels were increased.

Radford-Smith et al. (2022), in PND21 and PND112 mice, reported various alterations in metabolic markers in peripheral tissues. Blood and liver lactate levels increased at PND21, while blood lactate levels also increased at PND112. Conversely, alanine levels showed an increase in blood and liver at PND112 but decreased in blood at the same age, as well as in feces at PND112. Blood glucose levels decreased at both PND21 and PND112, while liver glucose levels increased at PND112. Taurine levels in the liver decreased at PND21 but increased at PND112. In fecal samples, ethanol exhibited a decrease at PND21 and an increase at PND112, alongside a decrease in alanine at PND21. Moreover, very low-density lipoprotein/triglycerides (VLDL/TG) and creatine levels decreased in blood and liver, respectively, at PND112. On the other hand, early prenatal *B. Dentium* did not affect blood indole levels at GD17.5 (Galley et al., 2024). When metabolic status of mice was evaluated with A. Muciniphila pre- and postantal treatment multiple changes in fecal metabolites were observed. At PND21, there were notable decreases in 2-deoxyadenosine, adenosine, phenylacetaldehyde, pyrimidine-derived compounds and trigonelline; while levels of 1,2,3-propanetricarboxylic acid, N-ethylglycine and L-carnitine increased. Finally, at PND42, a decrease was observed in L-lactic acid, N-ethylglycine, cytosine and tropine; whereas vitamin B7, indole-3-lactic acid, salicylic acid and N-desmethyltramadol levels increased (Qi et al., 2022).

### 3.7 Tight junction proteins and permeability

Only 3 studies with mice model analysed the impact in the permeability dependent on tight junction proteins. In brain PFC and gut, *B. Dentium*, *B. Bifidum* and *L. Lactis* prenatal supplementation result in an overexpressed *cldn3* in female and *cldn5* in male PND22-28 mice. In PND56-62 the treatment only increased *cldn3* expression in PFC of male mice. In gut tissue at this age, was *cldn5* mRNA levels in female mice the ones that have increased. These results found by Lathrop et al. (2024) in PFC and gut tissue are in line with the results obtained in gut by Zhou et al. (2022). *L. Rhamnosus* increased ZO-1 and CLDN3 protein concentration in PND98 mice and mRNA levels of *cldn3* in PND21 and PND98. In contrast, Qi et al. (2022) did not find effects in *zo1* or *cldn3* in PND21 and PND42 mice exposed pre- and postnatally exposed to *A. Muciniphila*. No effects were found in the gut levels of *ocldn* mRNA (Zhou et al., 2022; Qi et al., 2022) and only Lathrop et al. (2024) found increased *cldn1* expression in male treated mice.

### 3.8 Gut structure and function

6 of the 10 studies (40%) analysed the status of gut tissue in mice offspring. Regarding the proteins that play a key role in forming the protective mucus layer in the gut, Lathrop et al. (2024) found that *B. Dentium*, *B. Bifidum* and *L. Lactis* prenatal supplementation increased the expression of *muc2* and *muc3* in PND56-62 male mice.

*L. Rhamnosus* also increase the expression of *muc2* in PND21 and PND98 male and female mice (Zhou et al., 2022), however, Qi et al. (2022) did not find effects in *muc2* in young mice using *A. Muciniphila*. There is not too much information in the included articles about enteric nervous system related molecules, but no effects were found in the levels of substance P in PND49 mice (Zhu et al., 2022) or *ngf* expression in PND24 and PND86 rat (Moussavi et al., 2023). The results about gut structure integrity were divergent. On the one hand, Abu et al. (2023) found no significant effect on epithelial damage and inflammation in PND21 male mice treated with multi-strain pre- and postnatal probiotic. In contrast, Zhou et al. (2022) reported an increase in Ki67 positive cells, PAS positive cells, MUC2 positive cells, small IgA positive cells, and membrane localization of ZO-1 in PND21 mice exposed to *L. Rhamnosus*. Furthermore, an increase in 5-HT positive cells was observed in PND98 mice, while CagA positive cells showed no effect. Lastly, villi length and crypt depth increased in PND21 mice, but no significant changes were noted in PND98 mice, according to Zhou et al. (2022). These alterations of crypt and villi were not reported in PND21 mice when prenatal supplementation is composed of *L. Rhamnosus* (Qi et al., 2022).

### 3.9 Developmental effects

In the reviewed studies, several outcomes related to general development have been considered and represented in Table 5. No effects were observed on body weight across multiple studies involving both rat and mouse models, with findings spanning different PNDs during early development and adolescence (Xiao et al., 2020; Azagra-Boronat et al., 2016; Kim and Lee, 2022; Zhou et al., 2022; Lathrop et al., 2024; Qi et al., 2022). Interestingly, there were instances where body weight increased after pre- and postnatal multi-strain probiotic and *L. Rhamnosus* supplementation in mice at early stages of development between PND1 and PND28 (Radford-Smith et al., 2022; Zhou et al., 2022). In terms of brain measurements, both brain weight and length exhibited no significant changes (Azagra-Boronat et al., 2016). Similarly, gut weight and length did not demonstrate any notable effects (Azagra-Boronat et al., 2016). Lastly, the assessment of live births after probiotic gestational treatment indicated no discernible effect (Abu et al., 2023).

**Table 5.**
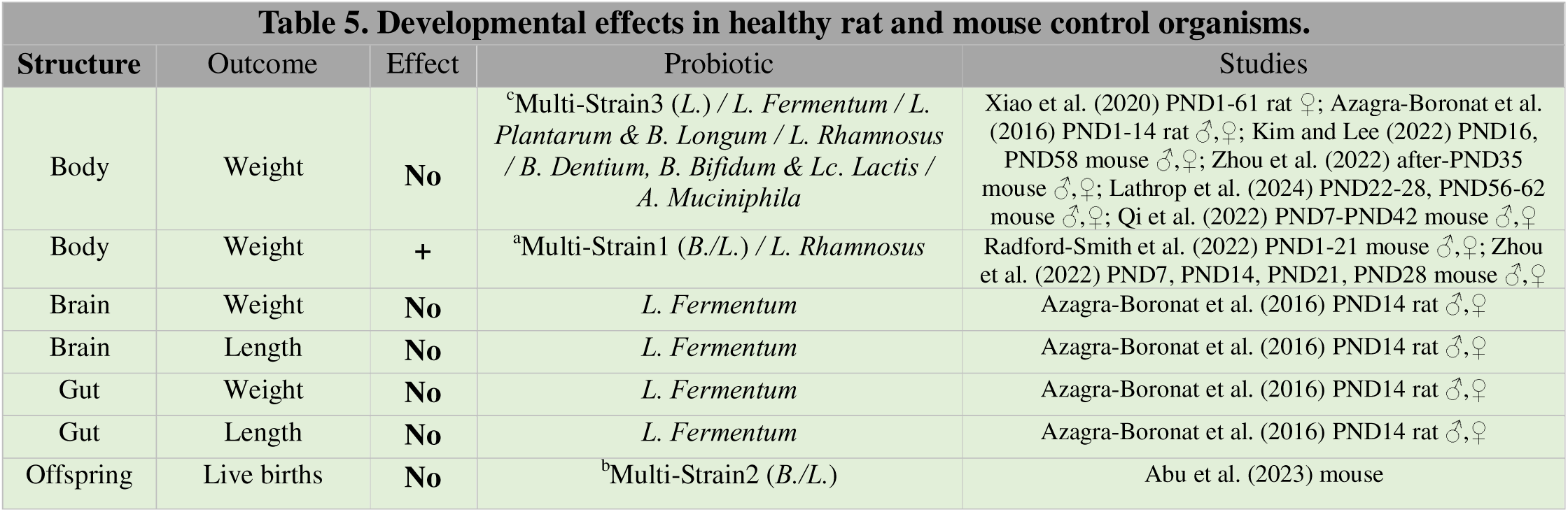
Developmental effects in healthy rat and mouse control organisms.

Abbr. aMulti-Strain1: B. Subtilis, B. Bifidum, B. Breve, B. Infantis, B. Longum, L. Acidophilus, L. Bulgaricus, L. Casei, L. Plantarum, L. Rhamnosus, L. Helveticus, L. Salivarius, Lc. Lactis and S. Thermophiles; bMulti-Strain2: L. Acidophilus, L. Bulgaricus, L. Paracasei, L. Plantarum, B. Breve, B. Infantis, B. Longum and S. Thermophilus; cMulti-Strain3: B. Longum, L. Acidophilus, L. Fermentum, L. Helveticus, L. Paracasei, L. Rhamnosus and S. Thermophilus; dMixture: B. Bifidum, B. Infantis, L. Helveticus, FOS and Maltodextrin.

### 3.10 Behavioral effects

A total of 12 of the studies (80%) evaluated some behavioral dimension in both mouse and rat models. The results were classified in Table 6.

**Table 6.**
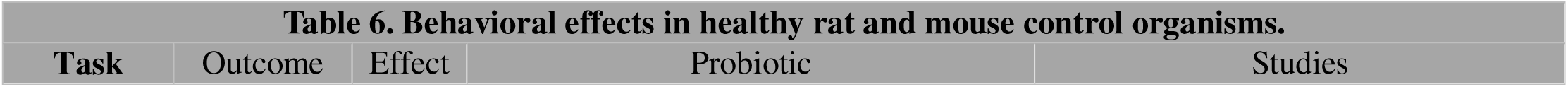
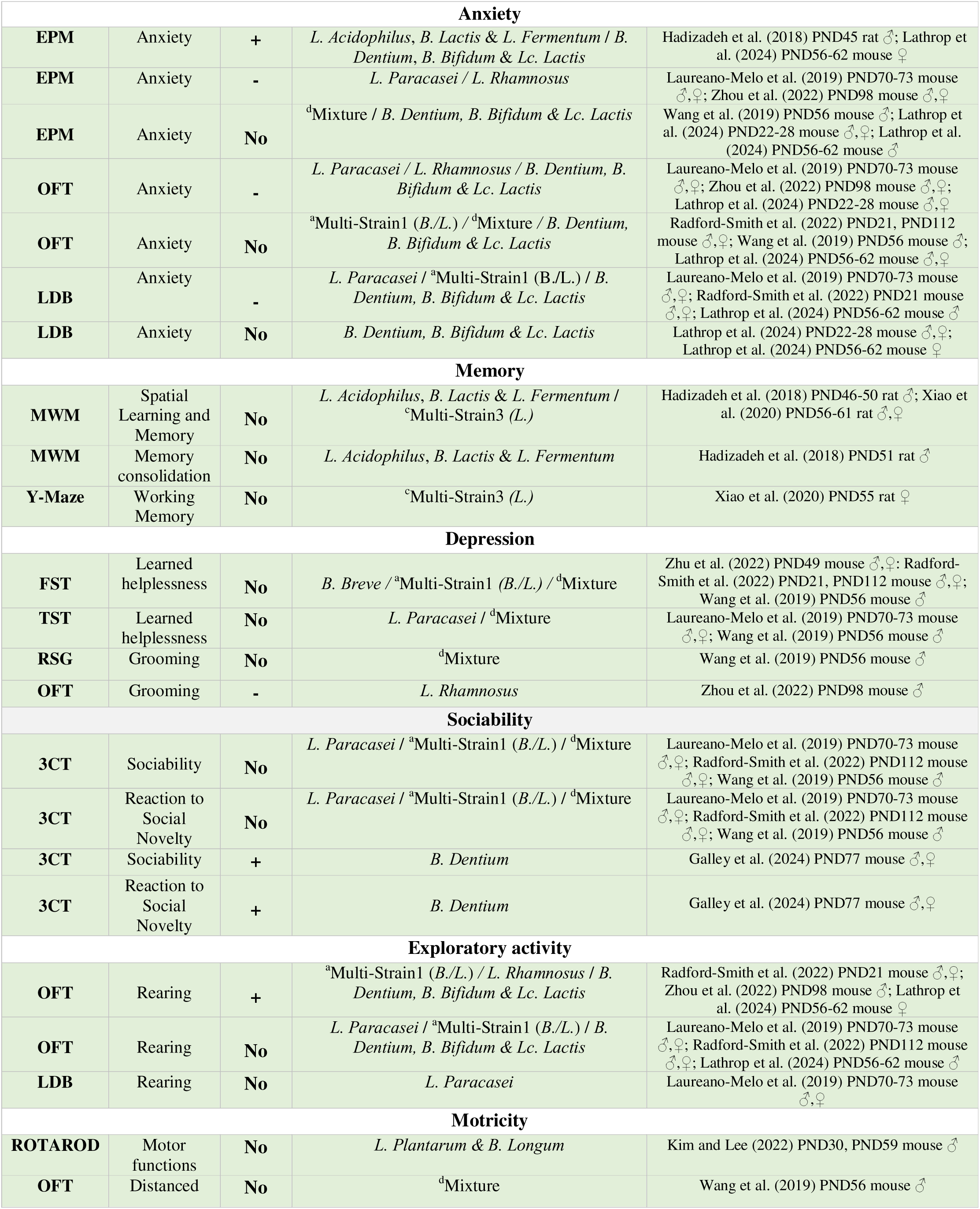
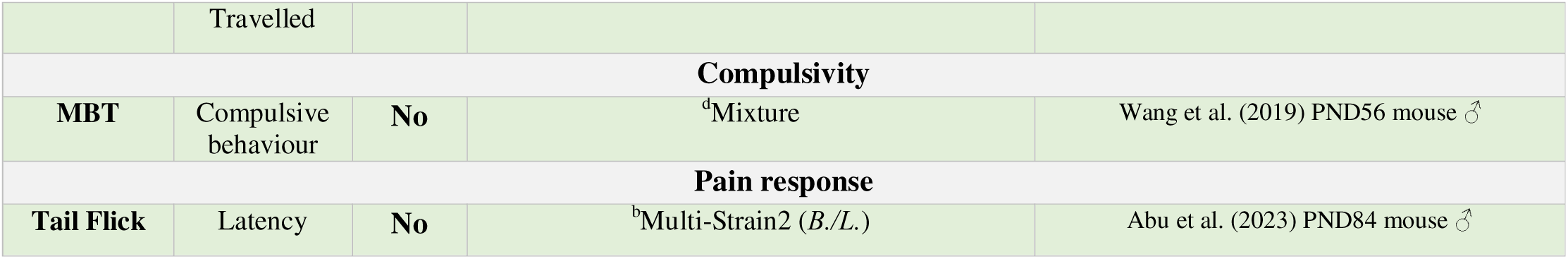
Behavioral effects in healthy rat and mouse control organisms.

| Task | Outcome | Effect | Probiotic | Studies |
| --- | --- | --- | --- | --- |

| Anxiety |  |  |  |  |
| --- | --- | --- | --- | --- |
| **EPM** | Anxiety | + | *L. Acidophilus, B. Lactis & L. Fermentum / B. Dentium, B. Bifidum & Lc. Lactis* | Hadizadeh et al. (2018) PND45 rat ♂; Lathrop et al. (2024) PND56-62 mouse ♀ |
| **EPM** | Anxiety | - | *L. Paracasei / L. Rhamnosus* | Laureano-Melo et al. (2019) PND70-73 mouse ♂,♀; Zhou et al. (2022) PND98 mouse ♂,♀ |
| **EPM** | Anxiety | **No** | <sup>d</sup>Mixture / *B. Dentium, B. Bifidum & Lc. Lactis* | Wang et al. (2019) PND56 mouse ♂; Lathrop et al. (2024) PND22-28 mouse ♂,♀; Lathrop et al. (2024) PND56-62 mouse ♂ |
| **OFT** | Anxiety | - | *L. Paracasei / L. Rhamnosus / B. Dentium, B. Bifidum & Lc. Lactis* | Laureano-Melo et al. (2019) PND70-73 mouse ♂,♀; Zhou et al. (2022) PND98 mouse ♂,♀; Lathrop et al. (2024) PND22-28 mouse ♂,♀ |
| **OFT** | Anxiety | **No** | <sup>a</sup>Multi-Strain1 (*B./L.*) / <sup>d</sup>Mixture / *B. Dentium, B. Bifidum & Lc. Lactis* | Radford-Smith et al. (2022) PND21, PND112 mouse ♂,♀; Wang et al. (2019) PND56 mouse ♂; Lathrop et al. (2024) PND56-62 mouse ♂,♀ |
| **LDB** | Anxiety | - | *L. Paracasei / <sup>a</sup>Multi-Strain1 (*B./L.*) / B. Dentium, B. Bifidum & Lc. Lactis* | Laureano-Melo et al. (2019) PND70-73 mouse ♂,♀; Radford-Smith et al. (2022) PND21 mouse ♂,♀; Lathrop et al. (2024) PND56-62 mouse ♂ |
| **LDB** | Anxiety | **No** | *B. Dentium, B. Bifidum & Lc. Lactis* | Lathrop et al. (2024) PND22-28 mouse ♂,♀; Lathrop et al. (2024) PND56-62 mouse ♀ |
| Memory |  |  |  |  |
| **MWM** | Spatial Learning and Memory | **No** | *L. Acidophilus, B. Lactis & L. Fermentum / <sup>c</sup>Multi-Strain3 (*L.*)* | Hadizadeh et al. (2018) PND46-50 rat ♂; Xiao et al. (2020) PND56-61 rat ♂,♀ |
| **MWM** | Memory consolidation | **No** | *L. Acidophilus, B. Lactis & L. Fermentum* | Hadizadeh et al. (2018) PND51 rat ♂ |
| **Y-Maze** | Working Memory | **No** | <sup>c</sup>Multi-Strain3 (*L.*) | Xiao et al. (2020) PND55 rat ♀ |
| Depression |  |  |  |  |
| **FST** | Learned helplessness | **No** | *B. Breve / <sup>a</sup>Multi-Strain1 (*B./L.*) / <sup>d</sup>Mixture* | Zhu et al. (2022) PND49 mouse ♂,♀; Radford-Smith et al. (2022) PND21, PND112 mouse ♂,♀; Wang et al. (2019) PND56 mouse ♂ |
| **TST** | Learned helplessness | **No** | *L. Paracasei / <sup>d</sup>Mixture* | Laureano-Melo et al. (2019) PND70-73 mouse ♂,♀; Wang et al. (2019) PND56 mouse ♂ |
| **RSG** | Grooming | **No** | <sup>d</sup>Mixture | Wang et al. (2019) PND56 mouse ♂ |
| **OFT** | Grooming | - | *L. Rhamnosus* | Zhou et al. (2022) PND98 mouse ♂ |
| Sociability |  |  |  |  |
| **3CT** | Sociability | **No** | *L. Paracasei / <sup>a</sup>Multi-Strain1 (*B./L.*) / <sup>d</sup>Mixture* | Laureano-Melo et al. (2019) PND70-73 mouse ♂,♀; Radford-Smith et al. (2022) PND112 mouse ♂,♀; Wang et al. (2019) PND56 mouse ♂ |
| **3CT** | Reaction to Social Novelty | **No** | *L. Paracasei / <sup>a</sup>Multi-Strain1 (*B./L.*) / <sup>d</sup>Mixture* | Laureano-Melo et al. (2019) PND70-73 mouse ♂,♀; Radford-Smith et al. (2022) PND112 mouse ♂,♀; Wang et al. (2019) PND56 mouse ♂ |
| **3CT** | Sociability | + | *B. Dentium* | Galley et al. (2024) PND77 mouse ♂,♀ |
| **3CT** | Reaction to Social Novelty | + | *B. Dentium* | Galley et al. (2024) PND77 mouse ♂,♀ |
| Exploratory activity |  |  |  |  |
| **OFT** | Rearing | + | <sup>a</sup>Multi-Strain1 (*B./L.*) / *L. Rhamnosus / B. Dentium, B. Bifidum & Lc. Lactis* | Radford-Smith et al. (2022) PND21 mouse ♂,♀; Zhou et al. (2022) PND98 mouse ♂; Lathrop et al. (2024) PND56-62 mouse ♀ |
| **OFT** | Rearing | **No** | *L. Paracasei / <sup>a</sup>Multi-Strain1 (*B./L.*) / B. Dentium, B. Bifidum & Lc. Lactis* | Laureano-Melo et al. (2019) PND70-73 mouse ♂,♀; Radford-Smith et al. (2022) PND112 mouse ♂,♀; Lathrop et al. (2024) PND56-62 mouse ♂ |
| **LDB** | Rearing | **No** | *L. Paracasei* | Laureano-Melo et al. (2019) PND70-73 mouse ♂,♀ |
| Motricity |  |  |  |  |
| **ROTAROD** | Motor functions | **No** | *L. Plantarum & B. Longum* | Kim and Lee (2022) PND30, PND59 mouse ♂ |
| **OFT** | Distanced | **No** | <sup>d</sup>Mixture | Wang et al. (2019) PND56 mouse ♂ |
|  | Travelled |  |  |  |
| **Compulsivity** |  |  |  |  |
| **MBT** | Compulsive behaviour | **No** | <sup>d</sup>Mixture | Wang et al. (2019) PND56 mouse ♂ |
| **Pain response** |  |  |  |  |
| **Tail Flick** | Latency | **No** | <sup>b</sup>Multi-Strain2 (*B./L.*) | Abu et al. (2023) PND84 mouse ♂ |

Abbr. ^a^Multi-Strain1: B. Subtilis, B. Bifidum, B. Breve, B. Infantis, 498 B. Longum, L. Acidophilus, L. Bulgaricus, L. Casei, L. Plantarum, L. Rhamnosus, L. Helveticus, L. Salivarius, Lc. Lactis and S.Thermophiles; bMulti-Strain2: L. Acidophilus, L. Bulgaricus, L. Paracasei, L. Plantarum, B. Breve, B. Infantis, B. Longum and S. Thermophilus; cMulti-Strain3: B. Longum, L. Acidophilus, L. Fermentum, L. Helveticus, L. Paracasei, L. Rhamnosus and S. Thermophilus; dMixture: B. Bifidum, B. Infantis, L. Helveticus, FOS and Maltodextrin.

#### 3.10.1 Anxiety

Anxiety was evaluated by 6 studies in rats and mice offspring using the light-dark box test (LDB), open field test (OFT), and plus maze test (EPM) paradigms. In LDB, the 3 studios found a decrease in anxiety levels in probiotic exposed mice. Supplementation with *L. Paracasei*, multi-strain probiotic (mainly *Lactobacillus* and *Bifidobacterium*) and *B. Dentium*, *B. Bifidum* and *L. Lactis* resulted in a increase in the time spent in the light compartment in PND70-73 mice, PND21 mice and PND56-62 male mice respectively (Laureano-Melo et al., 2019; Radford-Smith et al., 2022; Lathrop et al., 2024). However, no effects in anxiety measured by the LDB were found by Lathrop et al. (2024) in PND22-28 and PND56-62 female mice.

OFT also found decreases in anxiety in mice. Half of the studies (using *L. Paracasei*, *L. Rhamnosus* and *B. Dentium*, *B. Bifidum* and *L. Lactis*) found an increase in the time spent in center area in PND70-73, PND98 and PND22-28 male and female mice (Laureano-Melo et al., 2019; Zhou et al., 2022; Lathrop et al., 2024). Conversely, Lathrop et al. (2024) did not find effects in anxiety levels in PND56-62 mice of both sexes. Also, the other 2 studies did not observe altered OFT anxiety levels in the offspring exposed to multi-strain probiotic (PND21 and PND112 mice) (Radford-Smith et al., 2022) and in those treated with the mixture contained *B. bifidum, B. infantis, L. helveticus*, FOS and maltodextrin (PND56 male mice) (Wang et al., 2019).

Lastly, in EPM, more controversial results were found in anxiety levels measured by the time the animal spent in the open and closed arms. Hadizadeh et al. (2018) observed an increase in anxiety in PND45 male rats supplemented by *L. acidophilus, B. lactis* and *L. fermentum*, while Lathrop et al. (2024) found a similar increase in PND56-62 female mice when treatment was composed by *B. Dentium*, *B. Bifidum* and *L. Lactis*. In contrast, Laureano-Melo et al. (2019) pre- and postnatal *L. Paracasei* supplementation decreased anxiety in PND70-73 mice, as well as Zhou et al. (2022) in PND98 male mice (*L. Rhamnosus*). However, other studies, such as Wang et al. (2019) and Lathrop et al. (2024), found no significant effects on anxiety levels measured in EPM in PND56 male mice treated with the mixture contained *B. bifidum, B. infantis, L. helveticus*, FOS and maltodextrin; and PND22-28 mice and PND56-62 male mice.

#### 3.10.2 Depression-like behaviors

5 mice studies of the 15 total studies evaluated depressive-like behaviors in offspring prenatally supplemented with probiotics. No effects in immobility as an indicator of learned helplessness were found in forced swim test (FST) by Zhu et al. (2022), who focused on PND49 male and female mice prenatally treated with *B. Breve*; Radford- Smith et al. (2022), who examined PND21 and PND112 male and female mice supplemented by multi-strain probiotic; and Wang et al. (2019), who assessed immobility in PND56 male mice treated with *B. bifidum, B. infantis, L. helveticus*, FOS and maltodextrin. Immobility was also not altered in tail suspension test (TST) neither with pre- and postnatal *L. Paracasei* in PND70-73 mice nor with *B. bifidum, B. infantis, L. helveticus*, FOS and maltodextrin prenatal supplementation in PND56 male mice (Laureano-Melo et al., 2019; Wang et al., 2019). Lastly, grooming behavior was assessed in 2 studies only in male mice. While Wang et al. (2019) did not find effects in grooming in RSG test in PND56 male mice treated with the aforementioned mixture, Zhou et al. (2022) found that *L. Rhamnosus* decreased grooming evaluated in the context of OFT in PND98 male mice.

#### 3.10.3 Sociability

4 of the 15 studies explored how probiotic prenatal supplementation affects social behavior using the three-chamber test (3CT) only in mouse models. No effects were observed in general sociability and in reaction to social novelty in PND70-73 mice treated with pre- and postnatal *L. Paracasei*, PND112 mice supplemented with multi- strain probiotic and PND56 male mice prenatally supplemented with *B. Bifidum*, *B. Infantis*, *L. Helveticus*, FOS y maltodextrin (Laureano-Melo et al., 2019; Radford-Smith et al., 2022; Wang et al., 2019). In contrast, Galley et al. (2024) did indeed find an increase in both general sociability and reaction to social novelty in PND77, when mice were prenatally treated with *B. Dentium* during a brief period of gestation (GD10.5- GD16.5).

#### 3.10.4 Motricity and exploratory activity

4 mouse studies assessed rearing behavior in OFT, and a wide variety of results were obtained. While multi-strain pre- and postnatal probiotic increase rearing in PND21 mice, no effects in rearing were observed in adulthood (Radford-Smith et al., 2022). On the other hand, with *B. Dentium*, *B. Bifidum* and *L. Lactis* prenatal treatment, Lathrop et al. (2024) found a sexual dimorphism consisting of an increase in rearing only in PND56-62 female mice. In contrast, *L. Rhamnosus* increased rearing in OFT in PND98 male mice (Zhou et al., 2022). Lastly, *L. Paracasei* pre- and postnatal treatment did not affect rearing in PND70-73 mice of both sexes, neither in the OFT nor in the LDB (Laureano-Melo et al., 2019). Kim and Lee (2022) assessed motor functions with ROTAROD in male mice prenatally supplemented with *L. Plantarum* and *B. Longum* and no effects were found at PND30 and PND59 (Kim and Lee, 2022). Lastly distanced travelled in OFT was not affected in PND56 male mice in the research conducted by Wang et al. (2019).

#### 3.10.5 Other cognitive functions

In 2 studies that utilised rat models no effects were observed in spatial learning and memory consolidation assessed by Morris Water Maze (MWM) in PND46-51 male rat treated with *L. Acidophilus*, *B. Lactis* and *L. Fermuntum* until GD14; and PND56-61 rats of both sexes treated with multi-strain probiotic mainly composed of *Lactobacillus* strains (Hadizadeh et al., 2018; Xiao et al., 2020). No effects were observed in working memory assessed by Y-Maze in PND55 female rats (Xiao et al., 2020). Other cognitive functions found unaffected after probiotic prenatal treatment in male mice models were compulsivity assessed with marble burying test (MBT) in PND56 male mice (Wang et al., 2019) and pain response behavior in PND84 male mice (Abu et al., 2023).

### 3.11 Gut microbiota changes

**3.11.1 Actinobacteria**

All microbiota changes in offspring are depicted in Table 7. 46.7% of the studies did not examine how their probiotic treatments modify gut microbiota populations. The 50% of the studies that analyze the gut microbiota have found an increase in *Bifidobacterium* levels. Zhu et al. (2022) observed in both sexes an increase in *B. breve* in PND21 and PND42 mice following full-gestation exposure to this bacterial species and Abu et al. (2023) also reported an increase in PND21 mice with pre- and postnatal *Bifidobacterium*, *Lactobacillus* and *Streptococcus* exposure. On the other hand, with an exposure (GD18-PND5) to *L. rhamnosus*, this increase was observed in both male and female mice at PND98 (Zhou et al., 2022) Finally, in female rats, Xiao et al. (2020) also observed an increase in *Bifidobacterium* in PND68 after pre- and postnatal exposure to *Bifidobacterium*, *Lactobacillus* and *Streptococcus*.

**Table 7.**
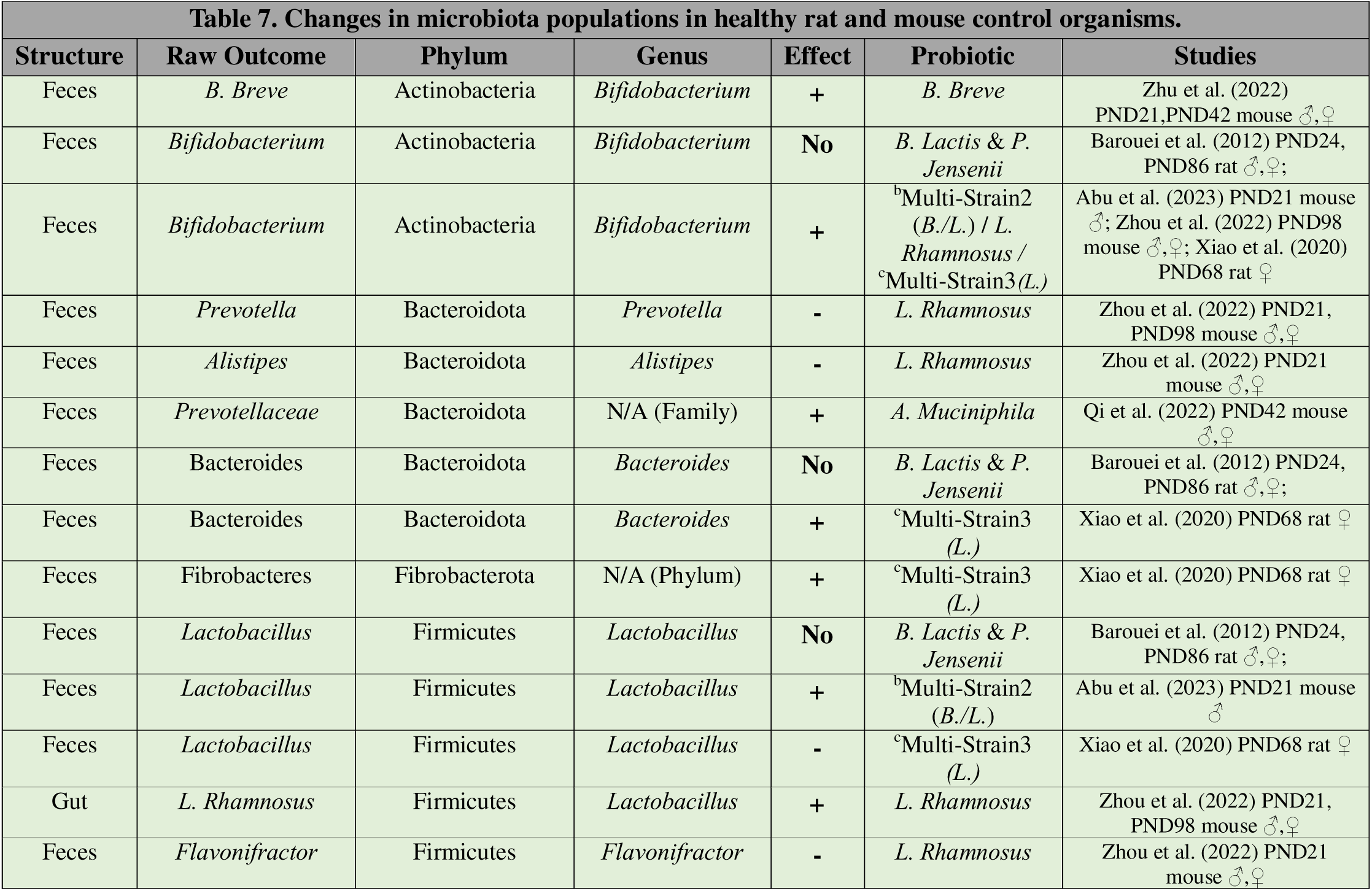
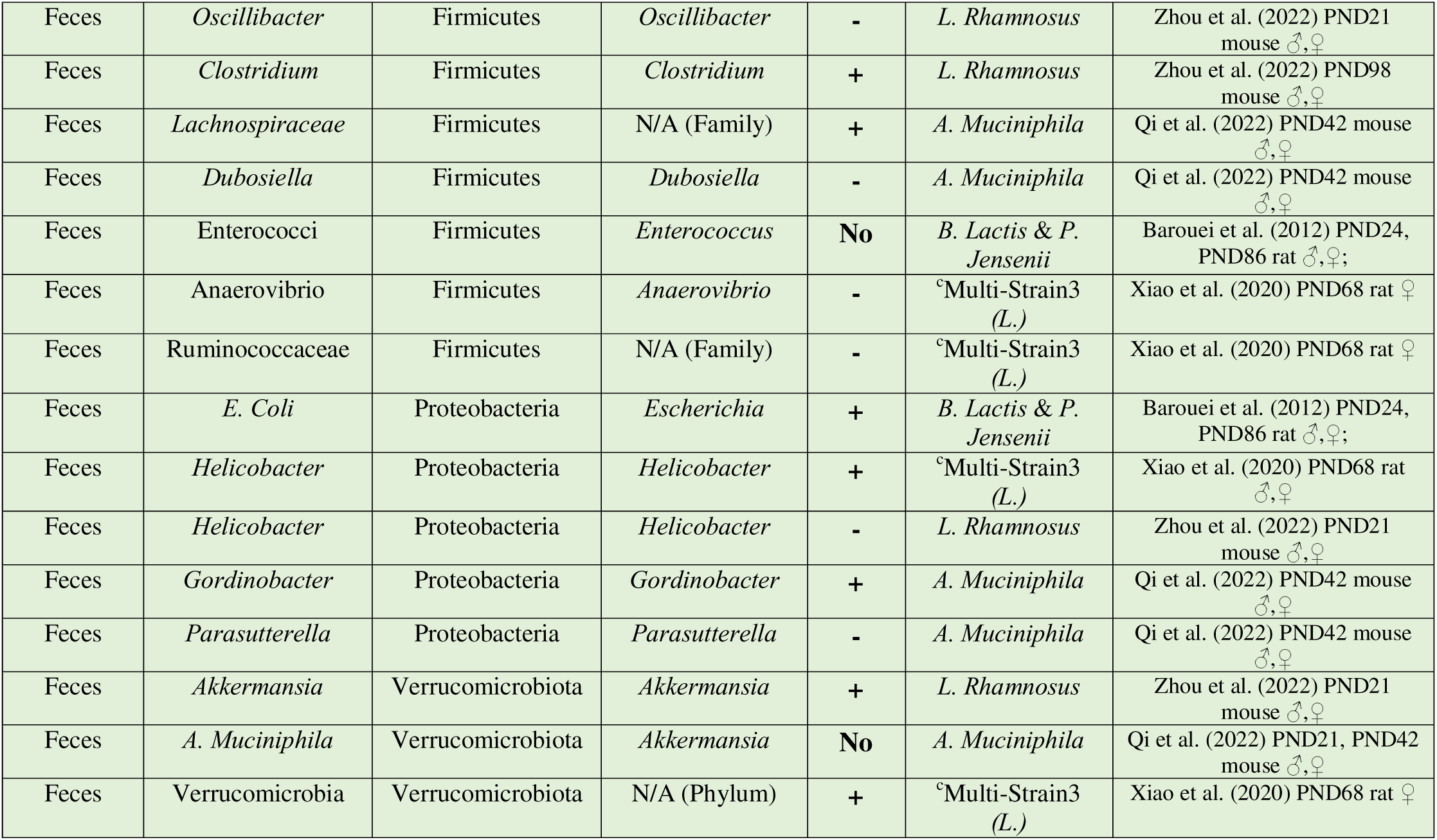
Changes in microbiota populations in healthy rat and mouse control organisms.

Abbr. ^a^Multi-Strain1: B. Subtilis, B. Bifidum, B. Breve, B. Infantis, B. Longum, L. Acidophilus, L. Bulgaricus, L. Casei, L. Plantarum, L. Rhamnosus, L. Helveticus, L. Salivarius, Lc. Lactis and S. Thermophiles; bMulti-Strain2: L. Acidophilus, L. Bulgaricus, L. Paracasei, L. Plantarum, B. Breve, B. Infantis, B. Longum and S. Thermophilus; cMulti-Strain3: B. Longum, L. Acidophilus, L. Fermentum, L. Helveticus, L. Paracasei, L. Rhamnosus and S. Thermophilus; dMixture: B. Bifidum, B. Infantis, L. Helveticus, FOS and Maltodextrin.

#### 3.11.2 Proteobacteria

62.5% of the studies found alterations in the *Proteobacteria* phylum, with 3 studies showing an increase in this phylum, 1 study showing a decrease, and the remaining study showing mixed results dependent on the genus. In rats pre- and postnatally exposed to the common genus *Lactobacillus*, Barouei et al. (2012) found an increase in *Escherichia* (*E. Coli*) levels in feces of PND24 and PND86 and Xiao et al. (2020) showed this increase in *Helicobacter* populations at PND68. In contrast, *L. Rhamnosus* treatment decreased *Helicobacter* levels in PND21 mice (Zhou et al., 2022). Finally, in PND42 mice when the probiotic is composed of *Akkermansia Muciniphila*, *Parasutterella* populations were decrease and *Gordinobacter* were increase (Qi et al., 2022).

#### 3.11.3 Firmicutes

A total of 50% studies showed differences in *Firmicutes* phylum. *Lactobacillus* pre- and postnatal treatment increased the levels of *Lactobacillus* in gut and feces of PND21 mice (Abu et al., 2023; Zhou et al., 2022). In addition, Zhou et al. (2022) found that *L. Rhamnosus* levels increased at PND98. In contrast, when this kind of treatment is proportionated to pregnant rats, the *Lactobacillus* populations were decreased in PND68 female rats (Xiao et al., 2020). Concerning the remaining genera within the *Firmicutes* phylum; *Lachnospiraceae* family, *Flavonifractor, Oscillibacter* and *Dubosiella* were decreased (Zhou et al., 2022; Qi et al., 2022) and *Clostridium* was increased in mice studies (Zhou et al., 2022). Finally, in PND68 rats principally treated with *Lactobacillus* we observed a decrease in *Ruminococcaceae* family and *Anaerovibrio* (Xiao et al., 2020).

#### 3.11.4 Bacteroidota, Fibrobacterota and Verrucomicrobiota

37.5% of studies found significant alterations in *Bacteroidota*. Xiao et al. (2020) found an increase in *Bacteroides* in PND68 rats. In mice, Zhou et al. (2022) showed a decrease in *Alistipes* and *Prevotella* after *L. Rhamnosus* supplementation, in contrast to the increase in the levels of *Prevotellaceae* populations with *Akkermansia Muciniphila* treatment (Qi et al., 2022). With respect to *Fibrobacterota*, only Xiao et al. (2020) found increase this phylum in PND68 rats.

Lastly, the study that supplemented *A. Muciniphila* found that this probiotic did not have effects in *A. Muciniphila* populations in mice at both ages PND21 and 42.

However, in *L. Rhamnosus* supplemented PND21 mice, we did observe an increase in *Akkermansia* genus. PND68 rats of Xiao et al. (2020) research had a general increase of *Verrucomicrobiota* phylum.

## 4. DISCUSSION

This review investigates the safety of prenatal probiotics, which are considered dietary supplements and do not undergo extensive testing like drugs. While often recommended during pregnancy to mitigate certain pathological risks, it is crucial to ensure their use does not negatively impact fetal development. The studies included in this review featured appropriate control groups, allowing us to group the main effects of gestational probiotics as reported in preclinical literature. However, they primarily employed probiotics with the objective of examining their effects within pathological models, such as autism models, *germ-free* rodents, prenatal high-fat diet, maternal stress, among others. Only Lathrop et al. (2024) specifically aimed to evaluate the safety of prenatal probiotics in relation to offspring health. The latter, using gestational *B. Dentium*, *B. Bifidum* and *Lc. Lactis*, is the only one who finds an up-regulation in oxytocin receptors in PFC accompanied by an increase in both general sociability and reaction to social novelty in 3CT. In contrast, other 3 studies which used other *Bifidobacterium* and *Lactobacillus* species did not find probiotics capable of alter sociability in adult mice (Laureano-Melo et al., 2019; Radford-Smith et al., 2022; Wang et al., 2019). Moreover, *Lactobacillus* did not impair memory or working memory in young rats (Hadizadeh et al., 2018; Xiao et al., 2020); however, it should be further investigated whether this effect could occur in late adulthood.

Several strains of *Lactobacillus* and *Bifidobacterium* were shown to influence anxiety- like behaviors in control organisms. For instance, studies employing the EPM, OFT and DLB tests demonstrated varying effects depending on the strain used and the timing of assessment postnatally. However, simple prenatal treatments with a single strain of *L. Paracasei* (Laureano-Melo et al., 2019) and *L. Rhamnosus* (Zhou et al., 2022) are shown to be stably able to reduce anxiety levels in different tasks when tested in adult mice, in contrast with the increase in anxiety levels performed in EPM in younger rats and mice prenatally exposed principally to *Bifidobacterium* genus and *L. Acidophilus* (Hadizadeh et al., 2018; Lathrop et al., 2024). Furthermore, brain tissue data suggest that GABAergic system could be more susceptible to *Lactobacillus* strain supplementations. In young mice, *Lactobacillus* genus have a significant impact on the early down-regulation of glutamic acid decarboxylase gene expression and low GABA levels in the whole brain (Radford-Smith et al., 2022; Laureano-Melo et al., 2019), while *Lactobacillus*-containing probiotics may promote GABA levels at later developmental stages (Radford-Smith et al., 2022; Zho et al., 2022). This biphasic effect on GABAergic system maturation could underlie the temporal differences in anxiety outcomes observed across studies. This aligns with the ability of *Lactobacillus* to produce GABA and to alter function at the central level in GABAergic neurons via the vagus nerve (Tette et al., 2022). Furthermore, in glutamatergic system following gestational administration of *Bifidobacterium* and *Lactobacillus*, we found an increase in glutamate levels in young mice (Radford-Smith et al., 2022; Kim and Lee, 2022), similar in age to those in studies where an apparently reduction in GABAergic function was found. Another evidence supporting the idea that these effects on excitatory/inhibitory balance seem to be more sensitive to the use of *Lactobacillus* strains is the finding by Wang et al. (2019), where no changes in glutamate levels were observed at the same age when primarily using *Bifidobacterium*. However, *Bifidobacterium* strains (especially *B. Lactis*) could influence the HPA axis and peripheral stress markers in both animal models (Barouei et al., 2012; Zhu et al., 2022; Moussavi et al., 2023). Further research with appropriate methods is needed to confirm these effects and to unravel the precise mechanisms through which gestational probiotics exert these effects to determine their long-term implications for mental health in healthy organisms.

On the other hand, gestational probiotics may have not a significant impact on the serotonergic system of control organisms. Outcomes about gestational *Bifidobacterium* and *Lactobacillus* influence in key genes related to 5-HT pathways (such as *tph2*, *5ht1a*, *5ht2a*, and *ido1*) have been found and no changes were observed across different brain regions and developmental stages in mice (Laureano-Melo et al., 2019; Galley et al., 2024; Radford-Smith et al., 2022). Also, no effects were found at peripheral level (Zhu et al., 2022; Zho et al., 2022; Galley et al., 2024). In fact, depression-like behaviors were unaffected with *Lactobacillus* and *Bifidobacterium* gestational supplementation (Zhu et al., 2022; Radford-Smith et al., 2022; Wang et al., 2019; Laureano-Melo et al., 2019). In contrast, Zhou et al. (2022) were the only authors that observed a detrimental in gut 5-HT function in adult mice with *L. Rhamnosus* and *B. Dentium*, and this impairment aligns with a reduction in grooming behavior in OFT. Unfortunately, this study did not explore its brain levels. To conclude, although actual literature shows that species such as *B. Longum* and *L. Rhamnosus* in postnatal probiotics have been able to affect the metabolism of 5-HT in gut and brain tissue (Li et al., 2019), prenatal treatments in healthy organisms did not affect significantly this system. However, further research is essential to determine whether prenatal administration of *L. rhamnosus* and *B. dentium* is truly safe for the development of the serotonergic system and mood regulation in adulthood, conclusion supported by the alterations found in Zhou et al. (2022), study that has a low risk of bias.

The alterations found in brain and peripheral metabolic status and fatty acids are anecdotal and totally dependent on methodological conditions of studies (n=1), so drawing general conclusions is unfeasible and a correct meta-analysis cannot be performed. Even so, it is important to highlight the large number of effects that have been found in healthy organisms (Azagra-Boronat et al., 2016; Qi et al., 2022; Radford- Smith et al., 2022; Zho et al., 2022; Zhu et al., 2022; Galley et al., 2024) and future lines of research should address this issue, being the effects of probiotics on fatty acids levels in pregnant women remain largely unexplored (Houttu et al., 2024). Surprisingly, the most of studies did not analyse the levels of SCFAs. Due to the production of SCFAs has proven to be one of the main mechanisms by which the microbiota is able to influence the CNS (Guo et al., 2022), more research about how gestational probiotics affects its levels is necessary. However, while prenatal *Bifidobacterium breve* supplementation did not affect total SCFAs in PND49 mice (Zhu et al., 2022), probiotics containing *Lactobacillus rhamnosus* and a mix of *Lactobacillus* and *Bifidobacterium* both altered peripheral acetate levels in young and adult mice (Radford-Smith et al., 2022; Zho et al., 2022). Acetate modulates immune and metabolic functions in microglia and can influence epigenetic regulation through histone and protein acetylation (Erny et al., 2021), effect that have been found in PND68 *Lactobacillus*-treated control rats (Xiao et al., 2019).

Regarding gut permeability, we can observe a tendency to increase tight junction proteins gene expression and protein levels with *L. Rhamnosus, B. Dentium, B. Bifidum* and *Lc. Lactis* (Zho et al., 2022; Qi et al., 2022; Lathrop et al., 2024). This effect of improvement of intestinal barrier function has been found previously with postnatal treatment in healthy intestinal epithelial cells with *L. Plantarum* in vitro treatment (Anderson et al., 2010). Furthermore, Lathrop et al. (2024) found that these strains of *Bifidobacterium* and *Lactococcus* also increased PFC levels of *cldn3* and *cldn5*, at the same time that increase the expression of the anti-inflammatory cytokine *il10* and *jam-a*. This last, encodes for a protein that regulates inflammation and blood-barrier functioning (Stamatovic et al., 2016). As lower permeability is associated with protection against pathological inflammatory responses (Anderson et al., 2010; Chen et al., 2017), these effects appear to be beneficial. However, small and larger molecules tight junction transport are a complex mechanism (see Zhao et al., 2022), and a disproportionately decrease in permeability could lead to difficulties in transport of ions and larger uncharged solutes (Milatz et al., 2010) that are crucial for maintaining ionic balance and membrane potential in cells.

Developmental effects have not been widely explored (such as eye opening, sexual development, amongst others) and, although 2 studies that used *L. Rhamnosus* in their supplementations found that its pre- and post-natal treatment increased body weight in the first four weeks of life in mice (Radford-Smith et al., 2022; Zho et al., 2022), probiotic gestational supplementation do not appear to be influencing the body weight evolution of offspring (Xiao et al., 2020; Azagra-Boronat et al., 2016; Kim & Lee, 2022; Zho et al., 2022; Lathrop et al., 2024; Qi et al., 2022). These results are consistent with those obtained by Wang et al. (2020) in their systematic review, concluding that prenatal probiotic supplementations in pregnant women with diabetes or overweight had minimal impact on newborn birth weight.

Based on the data about microbiota populations, probiotics seem to have different effects on various bacterial genera. For example, strains like *B. Breve* and multi-strain mixes that include *Lactobacillus* and *Bifidobacterium* lead to an increase in certain bacteria, especially *Bifidobacterium* and *Lactobacillus* (Zhu et al., 2022; Barouei et al., 2012; Abu et al., 2023; Zho et al., 2022; Xiao et al., 2020; Qi et al., 2022). On the other hand, *L. Rhamnosus* is often linked to changes in other genera, like *Prevotella*, *Alistipes*, and *Clostridium*, either increasing or decreasing their levels depending on the experimental conditions (Zho et al., 2022). Interestingly, some strains, such as *B. Lactis* and *P. Jensenii*, don’t seem to have a significant effect on certain bacterial populations such as bifidobacterium (Barouei et al., 2012), which suggests that not all probiotics have a broad-spectrum impact. Overall, it appears that multi-strain formulations and specific strains like *L. rhamnosus* tend to have a more noticeable effect on gut bacteria compared to single-strain probiotics. The increase in *Helicobacter* genus and *E. Coli* in several conditions (Barouei et al., 2012; Xiao et al., 2020) is particularly alarming because several *Helicobacter* species (Xu et al., 2022) and *E. Coli* (Schippa and Conte, 2014) could have harmful effects in health, and opportunistic infections have been proposed as one of the ways in which probiotics can damage host health (Merenstein et al., 2023).

Despite all the previously mentioned effects, they should be taken with caution due to the heterogeneity of the studies, which presents a significant challenge in drawing definitive conclusions about the effects of different bacterial genera administered during gestation on healthy subjects and controls in animal models. The small sample size of the outcomes obtained and the significant variations in methodologies used across studies complicate the ability to elucidate specific conclusions related to particular bacterial strains or doses. Regarding the latter, it is noteworthy that the doses used (ranging mostly from 10^6^ to 10^11^ CFU/day) are remarkably similar to those utilized in clinical studies investigating probiotics in humans, as well as to the concentrations of CFU/day found in probiotic supplements designed for human consumption—typically between 10^8^ and 10^11^ CFU/day (Ouwehand, 2017) —. Therefore, further research is necessary to determine the appropriate dosing in animal models to facilitate the extrapolation of results to human physiology.

To sum up, the strain-specific outcomes observed hinder the generalization of results, making it difficult to ensure the safety of prenatal probiotics use in healthy organisms without underlying pathologies. However, there is clear consensus that all studies reported effects on healthy control organisms, altering their physiology, and behavior in several cases, across different stages of life including infancy, adolescence and adulthood. Regarding the influence of the sex, there is apparently not enough evidence to point to a clear sexual dimorphism in the effects found. Moreover, probiotics, mainly *Lactobacillus* strains, are more sensitive in affecting GABAergic system maturation than serotonergic in healthy organisms, showing an apparent biphasic effect, reducing GABA functioning in early life but increasing them in later stages. Also, gestational probiotics can lead to gut dysbiosis in offspring and an over-increase in gut and brain permeability depending on tight junction proteins. This underscores the critical need to establish new lines of research employing appropriate methodologies to thoroughly investigate the safety of probiotic consumption during pregnancy, with particular attention to the effects of each bacterial species and the generation of more dedicated dose-response and security studies. Furthermore, this serves as a wake-up call to conduct similar investigations in healthy human populations, beyond the preclinical focus, to ensure that the findings are applicable and relevant to maternal and fetal health.

## DATA STATEMENT

The data supporting the findings of this study are available upon request. Researchers can access the data by contacting the corresponding author.

## REFERENCES

1. Abbasi, A., Aghebati-Maleki, A., Yousefi, M., & Aghebati-Maleki, L. (2021). Probiotic intervention as a potential therapeutic for managing gestational disorders and improving pregnancy outcomes. Journal of reproductive immunology, 143, 103244. 10.1016/j.jri.2020.103244

2. Abboud, M., Rizk, R., AlAnouti, F., Papandreou, D., Haidar, S., & Mahboub, N. (2020). The health effects of vitamin D and probiotic co-supplementation: A systematic review of randomized controlled trials. Nutrients, 13(1), 111.

3. Abu, Y. F., Singh, S., Tao, J., Chupikova, I., Singh, P., Meng, J., & Roy, S. (2024). Opioid-induced dysbiosis of maternal gut microbiota during gestation alters offspring gut microbiota and pain sensitivity. Gut microbes, 16(1), 2292224. 10.1080/19490976.2023.2292224

4. Anderson, R. C., Cookson, A. L., McNabb, W. C., Park, Z., McCann, M. J., Kelly, W. J., & Roy, N. C. (2010). Lactobacillus plantarum MB452 enhances the function of the intestinal barrier by increasing the expression levels of genes involved in tight junction formation. BMC microbiology, 10, 316. 10.1186/1471-2180-10-316

5. Azagra-Boronat, I., Tres, A., Massot-Cladera, M., Franch, À., Castell, M., Guardiola, F., Pérez-Cano, F. J., & Rodríguez-Lagunas, M. J. (2020). *Lactobacillus fermentum* CECT5716 Supplementation in Rats during Pregnancy and Lactation Impacts Maternal and Offspring Lipid Profile, Immune System and Microbiota. Cells, 9(3), 575. 10.3390/cells9030575

6. Barouei, J., Moussavi, M., & Hodgson, D. M. (2012). Effect of maternal probiotic intervention on HPA axis, immunity and gut microbiota in a rat model of irritable bowel syndrome. PloS one, 7(10), e46051. 10.1371/journal.pone.0046051

7. Bicknell, B., Liebert, A., Borody, T., Herkes, G., McLachlan, C., & Kiat, H. (2023). Neurodegenerative and Neurodevelopmental Diseases and the Gut-Brain Axis: The Potential of Therapeutic Targeting of the Microbiome. International journal of molecular sciences, 24(11), 9577. 10.3390/ijms24119577

8. Biosca-Brull, J., Pérez-Fernández, C., Mora, S., Carrillo, B., Pinos, H., Conejo, N. M., Collado, P., Arias, J. L., Martín-Sánchez, F., Sánchez-Santed, F., & Colomina, M. T. (2021). Relationship between autism spectrum disorder and pesticides: A systematic review of human and preclinical models. International Journal of Environmental Research and Public Health, 18(10), 5190. 10.3390/ijerph18105190

9. Cajal, B., Jiménez, R., Gervilla Garcia, E., & Montaño, J. J. (2020). Doing a systematic review in health sciences. Clínica y Salud, 31(2), 77–83. 10.5093/clysa2020a15

10. Chen, L., Deng, H., Cui, H., Fang, J., Zuo, Z., Deng, J., Li, Y., Wang, X., & Zhao, L. (2017). Inflammatory responses and inflammation-associated diseases in organs. Oncotarget, 9(6), 7204–7218. 10.18632/oncotarget.23208

11. Cuinat, C., Stinson, S. E., Ward, W. E., & Comelli, E. M. (2022). Maternal Intake of Probiotics to Program Offspring Health. Current nutrition reports, 11(4), 537–562. 10.1007/s13668-022-00429-w

12. Doi, M., Usui, N., & Shimada, S. (2022). Prenatal Environment and Neurodevelopmental Disorders. Frontiers in endocrinology, 13, 860110. 10.3389/fendo.2022.860110

13. Doron, S., & Snydman, D. R. (2015). Risk and safety of probiotics. Clinical Infectious Diseases, 60, S129–S134. 10.1093/cid/civ085

14. Erny, D., Dokalis, N., Mezö, C., Castoldi, A., Mossad, O., Staszewski, O., Frosch, M., Villa, M., Fuchs, V., Mayer, A., Neuber, J., Sosat, J., Tholen, S., Schilling, O., Vlachos, A., Blank, T., Gomez de Agüero, M., Macpherson, A. J., Pearce, E. J., & Prinz, M. (2021). Microbiota-derived acetate enables the metabolic fitness of the brain innate immune system during health and disease. Cell metabolism, 33(11), 2260–2276.e7. 10.1016/j.cmet.2021.10.010

15. Fijan S. (2014). Microorganisms with claimed probiotic properties: an overview of recent literature. International journal of environmental research and public health, 11(5), 4745–4767. 10.3390/ijerph110504745

16. Galley, J. D., King, M. K., Rajasekera, T. A., Batabyal, A., Woodke, S. T., & Gur, T. L. (2024). Gestational administration of Bifidobacterium dentium results in intergenerational modulation of inflammatory, metabolic, and social behavior. Brain, behavior, and immunity, 122, 44–57. 10.1016/j.bbi.2024.08.006

17. Guo, C., Huo, Y. J., Li, Y., Han, Y., & Zhouu, D. (2022). Gut-brain axis: Focus on gut metabolites short-chain fatty acids. World journal of clinical cases, 10(6), 1754– 1763. 10.12998/wjcc.v10.i6.1754

18. Hadizadeh, M., Hamidi, G. A., & Salami, M. (2019). Probiotic supplementation improves the cognitive function and the anxiety-like behaviors in the stressed rats. Iranian journal of basic medical sciences, 22(5), 506–514. 10.22038/ijbms.2019.33956.8078

19. Hooijmans, C. R., Rovers, M. M., de Vries, R. B., Leenaars, M., Ritskes-Hoitinga, M., & Langendam, M. W. (2014). SYRCLE’s risk of bias tool for animal studies. BMC medical research methodology, 14, 43. 10.1186/1471-2288-14-43

20. Houttu, N., Vahlberg, T., Miles, E. A., Calder, P. C., & Laitinen, K. (2024). The impact of fish oil and/or probiotics on serum fatty acids and the interaction with low-grade inflammation in pregnant women with overweight and obesity: secondary analysis of a randomised controlled trial. The British journal of nutrition, 131(2), 296–311. 10.1017/S0007114523001915

21. Kim, J., & Lee, S. (2022). Maternal low-intensity exercise and probiotic ingestion during pregnancy improve physical ability and brain function in offspring mice. Journal Of Functional Foods, 99, 105311. 10.1016/j.jff.2022.105311

22. Kolaček, S., Hojsak, I., Berni Canani, R., Guarino, A., Indrio, F., Orel, R., Pot, B., Shamir, R., Szajewska, H., Vandenplas, Y., van Goudoever, J., Weizman, Z., & ESPGHAN Working Group for Probiotics and Prebiotics (2017). Commercial Probiotic Products: A Call for Improved Quality Control. A Position Paper by the ESPGHAN Working Group for Probiotics and Prebiotics. Journal of pediatric gastroenterology and nutrition, 65(1), 117–124. 10.1097/MPG.0000000000001603

23. Laureano-Melo, R., Caldeira, R. F., Guerra, A. F., Da Conceição, R. R., De Souza, J. S., Giannocco, G., Marinho, B. G., Luchese, R. H., & Côrtes, W. S. (2019). Maternal supplementation with Lactobacillus paracasei DTA 83 alters emotional behavior in Swiss mice offspring. PharmaNutrition, 8, 100148. 10.1016/j.phanu.2019.100148

24. Lathrop, T., Perego, S., Bastiaanssen, T. F. S., van Hemert, S., Chronakis, I. S., & Diaz Heijtz, R. (2024). Multispecies probiotic intake during pregnancy modulates neurodevelopmental trajectories of offspring: Aiming towards precision microbial intervention. Brain, behavior, and immunity, 122, 547–554. 10.1016/j.bbi.2024.08.050

25. Li, H., Wang, P., Huang, L., Li, P., & Zhang, D. (2019). Effects of regulating gut microbiota on the serotonin metabolism in the chronic unpredictable mild stress rat model. Neurogastroenterology and motility, 31(10), e13677. 10.1111/nmo.13677

26. Merenstein, D., Pot, B., Leyer, G., Ouwehand, A. C., Preidis, G. A., Elkins, C. A., Hill, C., Lewis, Z. T., Shane, A. L., Zmora, N., Petrova, M. I., Collado, M. C., Morelli, L., Montoya, G. A., Szajewska, H., Tancredi, D. J., & Sanders, M. E. (2023). Emerging issues in probiotic safety: 2023 perspectives. Gut microbes, 15(1), 2185034. 10.1080/19490976.2023.2185034

27. Milatz, S., Krug, S. M., Rosenthal, R., Günzel, D., Müller, D., Schulzke, J. D., Amasheh, S., & Fromm, M. (2010). Claudin-3 acts as a sealing component of the tight junction for ions of either charge and uncharged solutes. Biochimica et biophysica acta, 1798(11), 2048–2057. 10.1016/j.bbamem.2010.07.014

28. Moussavi, M., Cuskelly, A., Jung, Y., Hodgson, D. M., & Barouei, J. (2023). Maternal probiotic intake attenuates ileal Crh receptor gene expression in maternally separated rat offspring. Bioscience, biotechnology, and biochemistry, 87(3), 308–313. 10.1093/bbb/zbac199

29. Radford-Smith, D. E., Probert, F., Burnet, P. W. J., & Anthony, D. C. (2022). Modifying the maternal microbiota alters the gut-brain metabolome and prevents emotional dysfunction in the adult offspring of obese dams. Proceedings of the National Academy of Sciences of the United States of America, 119(9), e2108581119. 10.1073/pnas.2108581119

30. Ouwehand A. C. (2017). A review of dose-responses of probiotics in human studies. Beneficial microbes, 8(2), 143–151. 10.3920/BM2016.0140

31. Ouzzani, M., Hammady, H., Fedorowicz, Z., & Elmagarmid, A. (2016). Rayyan-a web and mobile app for systematic reviews. Systematic reviews, 5(1), 210. 10.1186/s13643-016-0384-4

32. Qi, Y., Yu, L., Tian, F., Zhao, J., Zhang, H., Chen, W., & Zhai, Q. (2022). *A. muciniphila* Supplementation in Mice during Pregnancy and Lactation Affects the Maternal Intestinal Microenvironment. Nutrients, 14(2), 390. 10.3390/nu14020390

33. Sampson, T. R., & Mazmanian, S. K. (2015). Control of brain development, function, and behavior by the microbiome. Cell host & microbe, 17(5), 565–576. 10.1016/j.chom.2015.04.011

34. Sanders, M. E., Guarner, F., Guerrant, R., Holt, P. R., Quigley, E. M., Sartor, R. B., Sherman, P. M., & Mayer, E. A. (2013). An update on the use and investigation of probiotics in health and disease. Gut, 62(5), 787–796. 10.1136/gutjnl-2012-302504

35. Sanz, Y. (2011). Gut microbiota and probiotics in maternal and infant health. American Journal Of Clinical Nutrition, 94, S2000–S2005. 10.3945/ajcn.110.001172

36. Schippa, S., & Conte, M. P. (2014). Dysbiotic events in gut microbiota: impact on human health. Nutrients, 6(12), 5786–5805. 10.3390/nu6125786

37. Shamseer, L., Moher, D., Clarke, M., Ghersi, D., Liberati, A., Petticrew, M., Shekelle, P., Stewart, L. A., & PRISMA-P Group (2015). Preferred reporting items for systematic review and meta-analysis protocols (PRISMA-P) 2015: elaboration and explanation. BMJ (Clinical research ed.), 350, g7647. 10.1136/bmj.g7647

38. Sherwin, E., Bordenstein, S. R., Quinn, J. L., Dinan, T. G., & Cryan, J. F. (2019). Microbiota and the social brain. *Science (New York*, N.Y.), 366(6465), eaar2016. 10.1126/science.aar2016

39. Stamatovic, S. M., Johnson, A. M., Keep, R. F., & Andjelkovic, A. V. (2016). Junctional proteins of the blood-brain barrier: New insights into function and dysfunction. Tissue barriers, 4(1), e1154641. 10.1080/21688370.2016.1154641

40. Swartwout, B., & Luo, X. M. (2018). Implications of probiotics on the maternal- neonatal interface: Gut Microbiota, immunomodulation, and autoimmunity. Frontiers in Immunology, 9, 2840. 10.3389/fimmu.2018.02840

41. Tette, F. M., Kwofie, S. K., & Wilson, M. D. (2022). Therapeutic Anti-Depressant Potential of Microbial GABA Produced by *Lactobacillus rhamnosus* Strains for GABAergic Signaling Restoration and Inhibition of Addiction-Induced HPA Axis Hyperactivity. Current issues in molecular biology, 44(4), 1434–1451. 10.3390/cimb44040096

42. Vuong, H. E., & Hsiao, E. Y. (2017). Emerging Roles for the Gut Microbiome in Autism Spectrum Disorder. Biological psychiatry, 81(5), 411–423. 10.1016/j.biopsych.2016.08.024

43. Wang, C. C., Tung, Y. T., Chang, H. C., Lin, C. H., & Chen, Y. C. (2020). Effect of Probiotic Supplementation on Newborn Birth Weight for Mother with Gestational Diabetes Mellitus or Overweight/Obesity: A Systematic Review and Meta-Analysis. Nutrients, 12(11), 3477. 10.3390/nu12113477

44. Wang, X., Yang, J., Zhang, H., Yu, J., & Yao, Z. (2019). Oral probiotic administration during pregnancy prevents autism-related behaviors in offspring induced by maternal immune activation via anti-inflammation in mice. Autism research : official journal of the International Society for Autism Research, 12(4), 576–588. 10.1002/aur.2079

45. Wang, X., Yang, J., Zhang, H., Yu, J., & Yao, Z. (2019). Oral probiotic administration during pregnancy prevents autismCrelated behaviors in offspring induced by maternal immune activation via antiCinflammation in mice. Autism Research, 12(4), 576–588. 10.1002/aur.2079

46. Xiao, J., Wang, T., Xu, Y., Gu, X., Li, D., Niu, K., Wang, T., Zhao, J., Zhouu, R., & Wang, H. L. (2020). Long-term probiotic intervention mitigates memory dysfunction through a novel H3K27me3-based mechanism in lead-exposed rats. Translational psychiatry, 10(1), 25. 10.1038/s41398-020-0719-8

47. Xu, W., Xu, L., & Xu, C. (2022). Relationship between *Helicobacter pylori* infection and gastrointestinal microecology. Frontiers in cellular and infection microbiology, 12, 938608. 10.3389/fcimb.2022.938608

48. Yan, M., Man, S., Sun, B., Ma, L., Guo, L., Huang, L., & Gao, W. (2023). Gut liver brain axis in diseases: the implications for therapeutic interventions. Signal transduction and targeted therapy, 8(1), 443. 10.1038/s41392-023-01673-4

49. Zhao, Y., Gan, L., Ren, L., Lin, Y., Ma, C., & Lin, X. (2022). Factors influencing the blood-brain barrier permeability. Brain research, 1788, 147937. 10.1016/j.brainres.2022.147937

50. Zhou, B., Jin, G., Pang, X., Mo, Q., Bao, J., Liu, T., Wu, J., Xie, R., Liu, X., Liu, J., Yang, H., Xu, X., Wang, B., & Cao, H. (2022). Lactobacillus rhamnosus GG colonization in early life regulates gut-brain axis and relieves anxiety-like behavior in adulthood. Pharmacological research, 177, 106090. 10.1016/j.phrs.2022.106090

51. Zhu, H., Tian, P., Qian, X., Gu, L., Zhao, J., Wang, G., & Chen, W. (2022). Perinatal transmission of a probiotic *Bifidobacterium* strain protects against early life stress-induced mood and gastrointestinal motility disorders. Food & function, 13(14), 7520–7528. 10.1039/d2fo01164f

